# Directed evolution of colE1 plasmid replication compatibility: a fast tractable tunable model for investigating biological orthogonality

**DOI:** 10.1101/2021.11.25.470029

**Authors:** Santiago Chaillou, Eleftheria-Pinelopi Stamou, Leticia Torres, Ana B. Riesco, Warren Hazelton, Vitor B. Pinheiro

**Affiliations:** KU Leuven, Department of Pharmaceutical and Pharmacological Sciences, Rega Institute for Medical Research, Herestraat, 49, 3000, Belgium; University College London, Institute of Structural and Molecular Biology, University College London, Gower Street, London, WC1E 6BT, UK

**Author notes:** contributed equally to the publication (Santiago Chaillou, Eleftheria-Pinelopi Stamou).

**Keywords:** colE1 plasmids, plasmid orthogonality, origin of replication, plasmid copy number

## Abstract

Plasmids of the ColE1 family are among the most frequently used plasmids in molecular biology. They were adopted early in the field for many biotechnology applications, and as model systems to study plasmid biology. The mechanism of replication of ColE1 plasmids is well understood, involving the interaction between a plasmid-encoded sense-antisense gene pair (RNAI and RNAII). Because of its mechanism of replication, bacterial cells cannot maintain two different plasmids with the same origin, with one being rapidly lost from the population – a process known as plasmid incompatibility. While mutations in the regulatory genes RNAI and RNAII have been reported to make colE1 plasmids more compatible, there has been no attempt to engineer compatible colE1 origins, which can be used for multi-plasmid applications and that can bypass design constrains created by the current limited plasmid origin repertoire available. Here, we show that by targeting sequence diversity to the loop regions of RNAI (and RNAII), it is possible to select new viable colE1 origins that are compatible with the wild-type one. We demonstrate origin compatibility is not simply determined by sequence divergence in the loops, and that pairwise compatibility is not an accurate guide for higher order interactions. We identify potential principles to engineer plasmid copy number independently from other regulatory strategies and we propose plasmid compatibility as a tractable model to study biological orthogonality. New characterised plasmid origins increase flexibility and accessible complexity of design for challenging synthetic biology applications where biological circuits can be dispersed between multiple independent genetic elements.

## Introduction

Extrachromosomal genetic elements that replicate independently from the host’s own genetic information are common in nature. They have been reported in all of life’s kingdoms and encompass a diverse range of elements exploiting different topologies (linear or circular molecules), lengths (from kilo-to megabase-long molecules), copy number (from 1 to hundreds per cell), and reliant on different strategies for maintenance and replication within the host. Bacterial circular extrachromosomal double-stranded DNA genetic elements, better known as plasmids, have been instrumental for the development of molecular biology and remain one of its most common (and useful) tools. Within Synthetic Biology, significant standardization efforts have been made to streamline plasmids by sequence optimization (e.g. removal of non-functional sequences(1) and removal of unwanted endonuclease restriction sites(2)) and modularization – through academic (and commercial) initiatives such as the pSB1 plasmids used in iGEM, the Standard European Vector Architecture (SEVA)(3, 4), or even combinations thereof(5).

Simple applications, such as the heterologous expression of a single protein or the development of simple genetic circuits(6), can be readily carried out from a single plasmid without significant impact on plasmid purification, engineering and transformation. However, more complex applications such as the expression of large protein complexes(7, 8), optimization of synthetic pathways(9, 10), or directed evolution(11, 12) can benefit from the flexibility of having multiple plasmids within a single cell.

While it is possible in some cases to maintain multiple plasmids within a cell through continued selection, plasmids that share a common replication machinery tend to compete during replication, resulting (over the course of multiple generations) in a single plasmid being retained – the process commonly known as plasmid incompatibility(13, 14). Although as many as 27 incompatibility groups have been identified in the *Enterobactericeae* family(15), the number of replication mechanisms is significantly lower, demonstrating that replication orthogonality can be achieved, by means of sufficiently different replication machinery, even in situations where the replication strategy is shared. That is well established in molecular biology where some of the most commonly used plasmids have colE1-type origins but sit in different incompatibility groups: ColE1 (e.g. pUC, pET, pBR322, pMB1), p15A (e.g. pBAD, pACYC), CloDF13 (e.g. pCDF) and RSF1030 (e.g. pRSF)(16) – see **Figure 1**.

**Figure 1:**
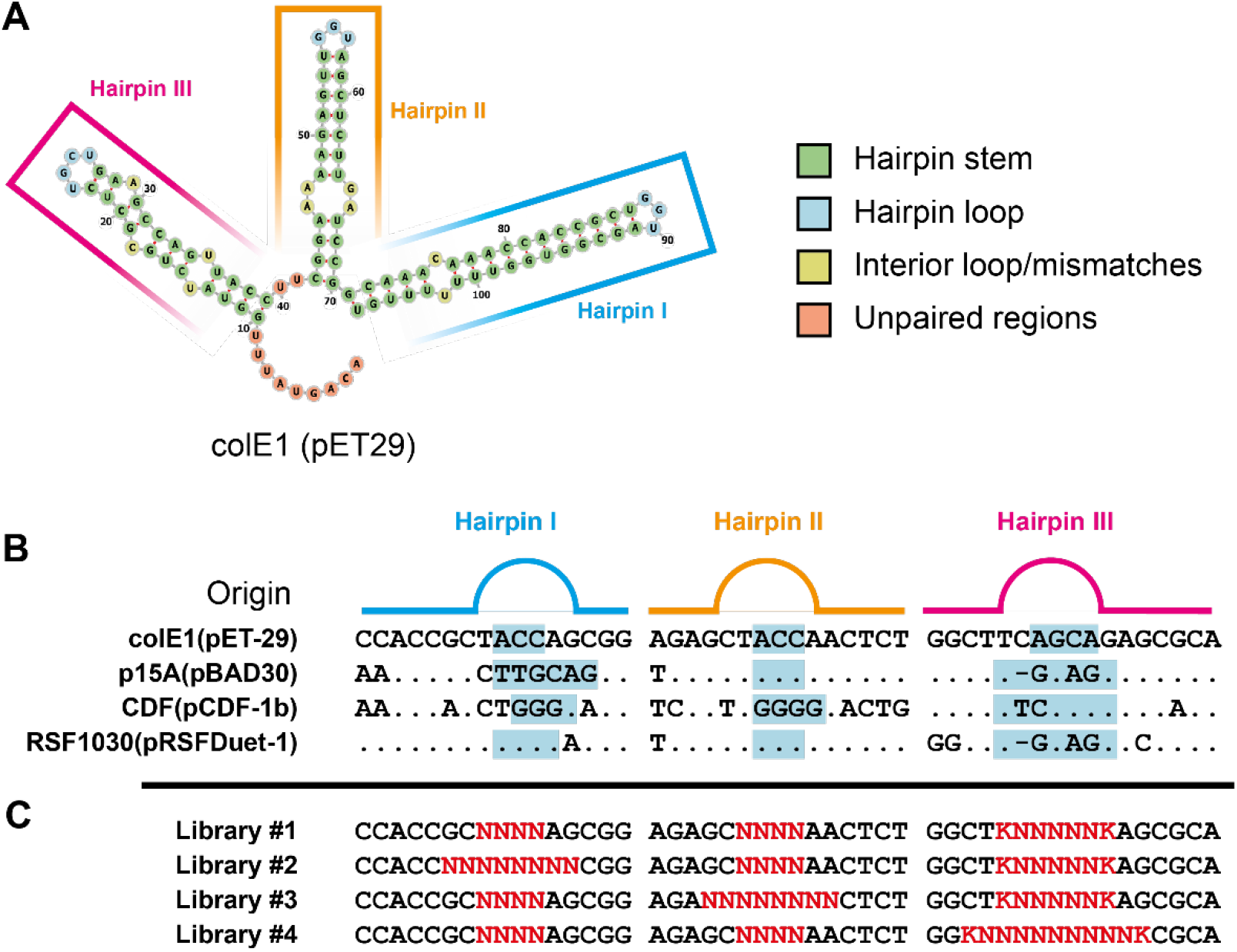
RNAI structure, sequence comparison between common colE1 origins and RNAI libraries. **A.** The predicted structure for RNA I, generated by the ViennaRNA suite(42), consists of three stem loops named (5’→3’) Hairpin III, II, and I. Nucleotides are coloured based on their position in the structure, whether in stem (green), loop (blue), stem bulges and mismatches (yellow), and unpaired (orange). **B.** Natural plasmids from different compatibility groups show similarities in sequence (shown as. in comparison to colE1 origin in pET29) in both stem and loops (highlighted in blue). **C.** Libraries were designed to introduce sequence variation (red) around loop regions. Sequence degeneracies are shown as per IUPAC (N = A, C, G or T; K = C, T).

Replication of colE1-type plasmids rely on the interplay between two RNA transcripts, from a sense-antisense overlapping gene pair (RNAI and RNAII), and the plasmid DNA(17, 18). The efficiency of those interactions has a direct impact on plasmid copy number per cell(19) and it is further modulated by plasmid-encoded dsRNA binding proteins such as Rom/Rop(20). Although RNAI and RNAII are complementary (and therefore their interaction is thermodynamically favourable), both molecules are highly structured (hence their interaction is not kinetically favourable)(21). Loop-loop interactions between RNAI and RNAII initiate and facilitate the formation of their dsRNA complexes, which as a result cannot initiate plasmid replication(19).

The well-characterised interactions between RNAI and RNAII is further supported by multiple investigations on the impact of mutations, insertions and deletions on the viability of the colE1 origin and on their potential effect on plasmid compatibility(18, 22–24). Together with the simplicity of readouts (presence or absence of a plasmid in a cell), colE1 origins of replications have the potential to be powerful model systems for studying biological orthogonality – how to engineer it, how to characterise it, and how to define it.

Orthogonality is a common metaphor in Synthetic Biology(25), representing a biological circuit or process that can operate independently and without interference from its natural equivalent(26, 27). In addition to its potential role in the biocontainment of engineered organisms(28), orthogonal systems can be easier to engineer (increased modularity) and control (fewer interactions with the natural cellular components). Directed evolution has been successfully applied for the isolation of orthogonal tRNA synthetases(29), bacterial two-component signalling systems(30) among other examples.

Here, we report the directed evolution and characterisation of multiple new colE1-type plasmid origins (compatible with the standard ColE1) based on diversification of the RNAI loops involved in the initial RNAI/RNAII interactions. We show that plasmid compatibility is a useful testbed for studying biological orthogonality, demonstrating that it is a continuum and that distance between states can be measured. We also identify possible routes towards the precise engineering of plasmid copy numbers.

## Material and Methods

### Bacterial strains, media, and culture conditions

*E. coli* DH5α (genotype: F^−^, ϕ80d/*αcZ*ΔM15, Δ(*lacZYA-argF*), U169, *end*A1, *hsd*R17(r_K_^−^, m_K_^+^), *rec*A1, *rel*A1, *pho*A, *sup*E44, *thi*-1, *gyr*A96, λ–) was used in all experiments. Cells were grown in Lysogeny broth (LB) medium (1% (w/v) tryptone, 0.5% (w/v) NaCl, 0.5% (w/v) yeast extract) unless otherwise stated. Liquid cultures were grown in aerated conditions (200 RPM) at 37°C, unless otherwise stated. For growth in solid medium, 1.5% (w/v) agar was added. Recovery after transformation was carried out in SOC Outgrowth Medium (New England Biolabs). The cell density of liquid cultures was estimated by measuring turbidity at 600 nm (OD600) in a SpectroStar Nano (BMG Labtech) spectrometer after blanking with fresh media.

When analysing the fluorescence of the co-transformed plasmids, several growth media were tested. These were LB/2 medium (0.5% (w/v) tryptone, 0.25% (w/v) NaCl, 0.25% (w/v) yeast extract), LB*2 medium (2% (w/v) tryptone, 1% (w/v) NaCl, 1% (w/v) yeast extract), M9 medium (Na_2_HPO_4_ 33.7 mM, KH_2_PO_4_ 22.0 mM, NaCl 8.55 mM, NH_4_Cl 9.35 mM, glucose 0.4 % (w/v), MgSO_4_ 1 mM, CaCl_2_ 0.3 mM, BME vitamins (Merck) 1X), and Mueller-Hinton (MH) medium (0.2 % (w/v) beef extract, 1.75 % (w/v) casein hydrolysate, 0.15 % (w/v) starch).

Kanamycin (50 μg/mL), Chloramphenicol (34 μg/mL) or Ampicillin (50 μg/mL) were used as selection markers. Antibiotic free-media were kept at room temperature, antibiotic stocks at −20°C and antibiotic-containing media at 4°C.

### Plasmids and libraries construction

The plasmids described in this work (sequences in the Supplementary Information) were assembled via Type IIS(31) (using BsaI or BsmBI) or Gibson(32) assembly, from PCR-amplified or commercially synthesised (IDT, Leuven, BE) double-stranded DNA fragments, and available vectors: pET-29a (Novagen, US), pSB1C3 (iGEM Foundation), and pBAD expressing fluorescent protein variants (Addgene, UK). Primers used in the assembly of the vectors are listed in **SI Table 1**.

RNAI libraries were commercially synthesised as single-stranded oligonucleotides (IDT, Leuven, BE) and PCR amplified to introduce BsaI sites used in the subsequent Type IIS cloning into a PCR-amplified pET29a backbone. Primers and PCR conditions are given in the Supplementary Information.

### Serial cultures for microbiological compatibility assay and selection

Single plasmid or co-transformed cells were grown overnight in 5 mL of LB (in a 50 mL centrifuge tube) with the appropriate antibiotic selection (none, kanamycin 50 μg/mL, chloramphenicol 34 μg/mL, or both). Fresh cultures were inoculated from the overnight cultures (1:100) and appropriate antibiotics to start the experiments. Large-scale cultures (200 mL) were grown in 1 L conical flasks. Small-scale cultures were grown in 200 μL in flat- or round-bottom 96-well plates. For serial culturing, passaging was done after 24 h growth with new cultures started with 1% (v/v) inocula. Serial culturing was carried out between four and seven days, as described in the text. For quantification, cultures were diluted in phosphate-buffered saline (Sigma-Aldrich, UK) and plated overnight in LB-agar containing no antibiotics (for total cell quantification), kanamycin (for isolating total cells containing the reporter plasmid) or both (for measuring cells that did not lose any plasmid). Resulting plates were photographed or scanned in a GE Healthcare Typhoon FLA9500 biomolecular imager, using FITC settings (Laser: 488 nm; filter BLP) to identify GFP-expressing colonies (where relevant), and Alexa Fluor 647 (Laser: 635 nm; filter LPR) to visualize all *E. coli* colonies.

Small-scale cultures, used in the high-throughput compatibility assays, were grown for 24 h before cell density and fluorescence were measured on a CLARIOstar spectrophotometer (BMG Labtech, DE), with the 525/25 filter for GFP expression quantification. Gain settings for fluorescence measurements were automatically selected by the software based on the available samples, to maximise discrimination without saturating detection. Some selected samples of small-scale cultures, together with positive and negative controls, were also analysed in flow cytometry experiments.

Selection for compatible plasmid origins was carried out in large-scale cultures (from which alpha origin was isolated) and small-scale high-throughput screening assay (from which the characterised origins were isolated). Selection was carried out as per serial passaging experiments isolating compatible origins from bacterial colonies grown in both kanamycin and chloramphenicol after a period of growth under kanamycin only selection.

### Intercompatibility assay

The co-transformed cells were grown overnight in 5 mL of LB with kanamycin (50 μg/mL), chloramphenicol (34 μg/mL) and ampicillin (50 μg/mL) for 16 hours. Small-scale fresh cultures were inoculated from the overnight ones [1:100 (v/v)] and grown overnight in aerated conditions (200 RPM) at 37°C in 96-well plates. Serial passaging was carried out as described above, with media supplemented with chloramphenicol, three times before being analysed by flow cytometry. Expression of fluorescent proteins was induced in the final overnight growth cycle by the addition of L-arabinose [0.5% (w/v)].

### DNA extraction and copy number calculation using dPCR

Both genomic and plasmid DNA were extracted from *E. coli* cells, using a detergent-based method previously described(33). The isolated DNA was restriction digested with XbaI (NEB), following manufacturer’s recommendations on reaction conditions, to improve the efficiency of digital PCR amplification(34). The digital PCRs (dPCRs) were carried out using a ThermoFisher QuantStudio 3D platform.

TaqMan assays(35) were used to detect both genome and regulatory plasmid DNA. The probe for genome detection was designed against the Ter region of the *E. coli* genome. Being the last part of the bacterial chromosome to replicate, the Ter region minimises the overestimate of *E. coli* genomes due to replicating genomic DNA. The probe for plasmid detection was designed against the chloramphenicol acetyltransferase gene. The sequences of the primers and the probes are listed in **SI Table 1**.

Reaction mixes were prepared in 15 μL, containing QuantStudio™ 3D Digital PCR Master Mix v2 (1x), PCR primers (500 nM), FAM probe (500 nM) for plasmid detection, HEX probe (620 nM) for genome detection, and template 0.2 pg/μL. Once assembled, 14.5 μL of the dPCR mix were used per chip. Negative controls without plasmids (i.e. genome only) or with purified plasmids (i.e. no genome) were also carried out. Reaction conditions consisted of an initial denaturation at 96°C for 10 minutes, followed by 39 cycles of 60°C for 2 minutes, and 98°C for 30 seconds. A final 60°C extension for 2 minutes was carried out as a polishing step.

The data analysis was done using the QuantStudio^®^ 3D AnalysisSuite™ online platform. Using the rare mutation analysis for the signals amplified from the genomic DNA, the software provides the values for the target/total signals with a 95% confidence interval. These values allowed us to calculate the copy number of the plasmids per every genomic sequence.

### Flow Cytometry

Experiments were designed to detect bacterial expression of fluorescent proteins in the cross-compatibility (GFP only) and intercompatibility (GFP, mApple and mTagBFP2) assays, using cells transformed with single (or two) plasmids as controls to establish gates and check compensation parameters. Experiments were carried out in a CytoFLEX (Beckman Coulter, USA) benchtop flow cytometer. Super-folder GFP (sfGFP) was measured using FITC settings (laser: 488 nm, filter 525/40 BP). The mApple protein could be detected using the PE settings (laser: 488 nm, filter 585/42 BP). mTagBFP2 was measured using PB450 settings (laser : 405 nm, filter 450/45 BP).

Small culture aliquots (5 μL) were diluted in 1000 μL of Mili-Q H_2_O, and those samples were used for sampling. Forward and side scattering (FSC and SSC) were used to identify single bacterial cells (from noise, cell debris and aggregates) and gated accordingly for acquisition. All subsequent analyses were carried out in the gated cells using CytExpert (Beckman Coulter) or FCS Express (De Novo Software, USA). Single-cell gating was repeated post-acquisition and samples analysed, using in all circumstances at least 1300 events – See **SI Table 2** for details.

### NGS data analysis

Transformed libraries (Library #1 for method development and combined libraries for subsequent experiments), were harvested from the transformation plates and grown in liquid LB medium (50 mL culture under aerated conditions) supplemented with kanamycin at 37°C for 2 hours or until turbid. Cells were isolated by centrifugation (4,000 × *g* at 4°C for 15 min) and plasmid DNA isolated using a GeneJet mini-prep kit (ThermoFisher, UK) following manufacturer’s recommendations. Isolated plasmid DNA was quantified by UV absorbance and used as template for the amplification of the fragment to be sequenced (primers given in Supplementary Information). Sample preparation was carried out according to the recommendations of Genewiz, who also carried out the Illumina sequencing.

Compatible libraries, obtained after selection in large-scale cultures and plated under both kanamycin and chloramphenicol selection, were treated as above, which also explains the high frequency of observation of the wild-type colE1 origin in the analysis (SI_Fig 3).

NGS data analysis was carried out in the Galaxy server at *usegalaxy.org*(36). The analysis workflow (provided as supplementary information) begins with filtering sequences for quality using the ‘FASTQ Quality Trimmer’ (10 window, mean score >20). Sequences are converted from FASTQ to FASTA (fastq.info) and filtered with sequences expected to flank the library diversity (Filter FASTA). In our analysis, “CGTAATCTGCTGCTTGCAAA”, “GGTTTGTTTGCCGGA”, “TTTCCGAAGGTAACT” and “AGCGCAG” were used to reduce the data to sequences likely to be from bonafide origins. As a last step, the dataset is de-duplicated and repeated origins counted (Unique.seqs).

### Bioinformatic tools for RNA structure prediction

Secondary structure prediction of wild-type and isolated RNAI was obtained using the Vienna RNA RNAfold online server(37, 38), and visualised using forna(39).

## Results and Discussion

Engineering orthogonal biological components poses always a two-fold problem: specificity needs to be built upon activity, i.e. orthogonal origins of replication need to be functional (capable of supporting stable plasmid replication and maintenance). Therefore, we first focused in generating novel RNAI molecules that could support plasmid replication.

### Library design and selection of viable ColE1 variants

Directed evolution is a powerful approach for engineering biological components but considerations over sampling and density of the sequence landscape need to be taken into account(40). Although RNAI is the shorter RNA regulator, its length still exceeds 100 bases, thus thorough sampling of its available sequence space (1.6 × 10^60^ variants) is not possible. RNAII structure is known to be an integral part of its mechanism(41), therefore we reasoned that targeting sequence variation to the hairpins’ stems would likely disrupt their structure, creating a sparsely populated sequence space – even if alternative stems could lead to greater orthogonality.

Restricting variation to the loops (and their immediate vicinity) limits the maximum size of the search space (4.3 × 10^9^ variants) and would be expected to minimise disruption of the ternary structure of the hairpins. DNA oligonucleotides containing the required degeneracy (Figure 1) were used to generate origin libraries by PCR and subsequent Type IIS cloning. Transformation of the assembled libraries served therefore as a selection mechanism to isolate viable plasmids.

Viable plasmids isolated from the transformation of Library #1, were recovered and sampled by NGS (**SI Fig 1** and **SI Table 3**). It confirmed that alternative loops are possible and that all loops can tolerate diversification. Notably, whether artefacts of NGS sample preparation, or active *in vivo* selection during recovery, deletions and insertions were also observed both within and outside of the expected diversity – with a potential hotspot near the 3’-end of stem IIIb (**SI Fig 1**).

### Selecting colE1 compatible origins by serial passaging

Given that single point mutations in RNAI have been shown to be sufficient to affect plasmid incompatibility(18, 23), we co-transformed viable Library #1 plasmids into cells harbouring pSB1C3 (containing a wild-type colE1 origin) to investigate how best compatible origins could be isolated.

Repeated passaging of a bacterial culture in the absence of antibiotic selection is the traditional microbiological route towards quantifying plasmid stability and compatibility. As a selection tool, the simple readout (bacterial growth) and the experimental flexibility (media composition) make serial passaging a powerful strategy – capable of modifying even core biological processes such as the genetic code(43).

Our initial results confirmed that it could be used in plasmid compatibility selection, with 1% of the population still harbouring two plasmids after six continuous days of growth (data not shown). To increase selection pressure (towards loss of the variants) and minimise potential sources of external contamination, we repeated the serial passaging selection in the presence of chloramphenicol (thus forcing maintenance of the wild-type colE1 origin in pSB1C3). After two passages (approx. 80 generations), most of the cells still harboured two plasmids (**SI Fig. 2**).

Some of the colonies isolated from the selection for compatibility were sequenced, with one also being identified as the most frequent compatible variant by NGS (**SI Fig. 3** and **SI Table 4**). The variant, termed alpha (**SI Fig. 1B**), was regrown under compatibility selection conditions and compared to a wild-type colE1 origin. As expected, differences in compatibility were already observable after a single day growth and the alpha origin quantitatively remained in the population throughout the 4-day experiment (**Figure 2**).

**Figure 2:**
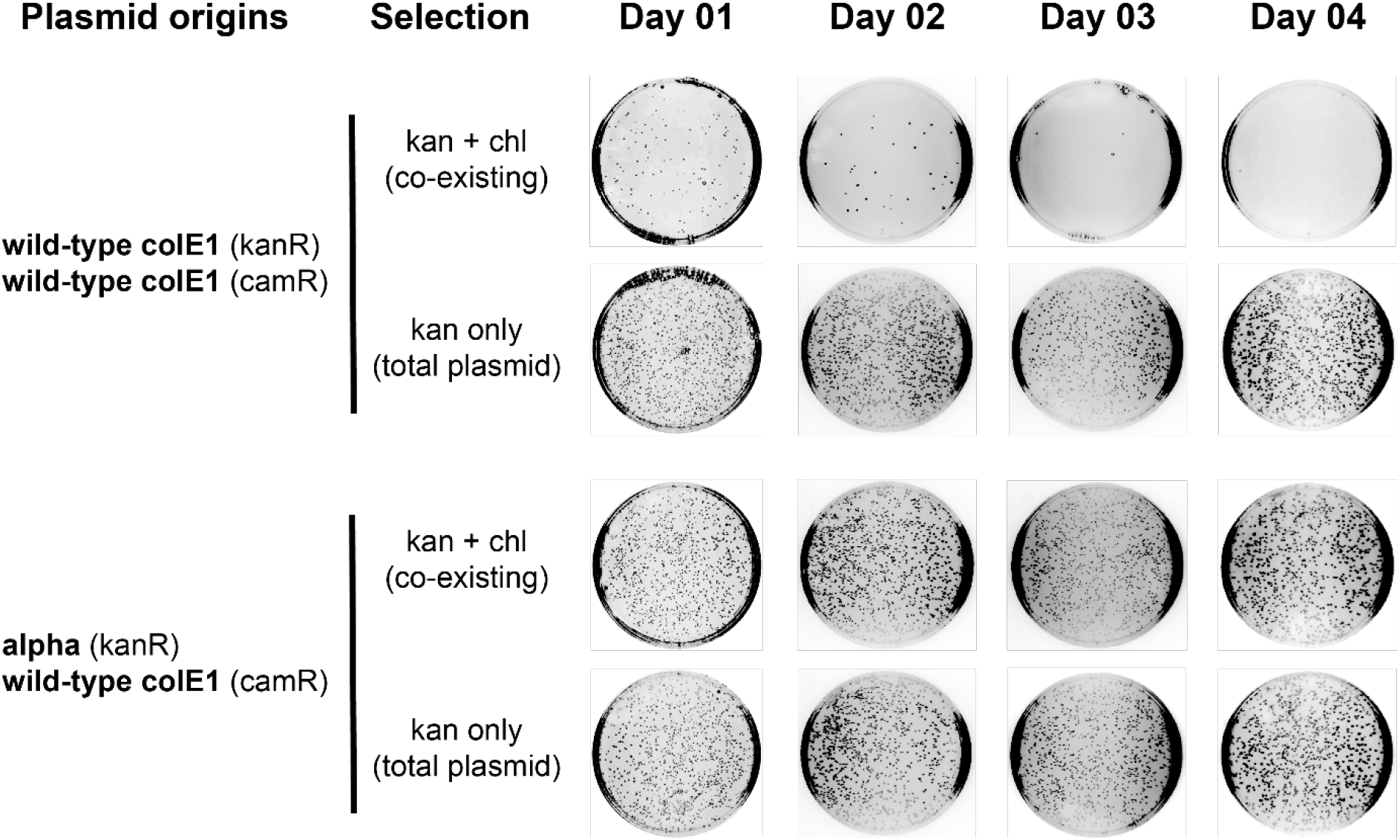
Microbiological plasmid compatibility assay. E coli cells harbouring two plasmids (maintained by antibiotic co-selection) are cultured only in the presence of kanamycin allowing the maintenance of the second plasmid (harbouring chloramphenicol resistance marker) to be tested. Aliquots from the cultures were diluted and plated in single (kanamycin) or double antibiotic plates, enabling estimates of plasmid maintenance to be calculated. While the second wild-type origin is rapidly lost (top), the wild-type origin co-cultured with the selected alpha origin (bottom) stably remains in the culture.

Although powerful as a selection strategy, serial passaging lacks an assay to report on the selection progress, and it remains a low-throughput and labour-intensive screening tool. We therefore chose to explore possible biological circuits that could be used as reporters in selection and as high-throughput screening to assess plasmid compatibility.

### High-throughput plasmid compatibility assays and selection

Fluorescent or coloured proteins are convenient and widely used reporters in bacteria, but their long half-lives and potential toxicity(44, 45) can limit their usefulness in dynamic processes and also lead to an increased metabolic burden on the cells – in this case potentially exacerbating the stress cells may already be under from harbouring multiple colE1 high copy number plasmids. In addition, metabolic burden and protein expression are both known factors in colE1 plasmid copy number variability(46, 47). Therefore, we opted for the biological circuit shown in **Figure 3**, in which a TetR-based negative feedback loop minimises any potential metabolic burden from reporter expression.

**Figure 3:**
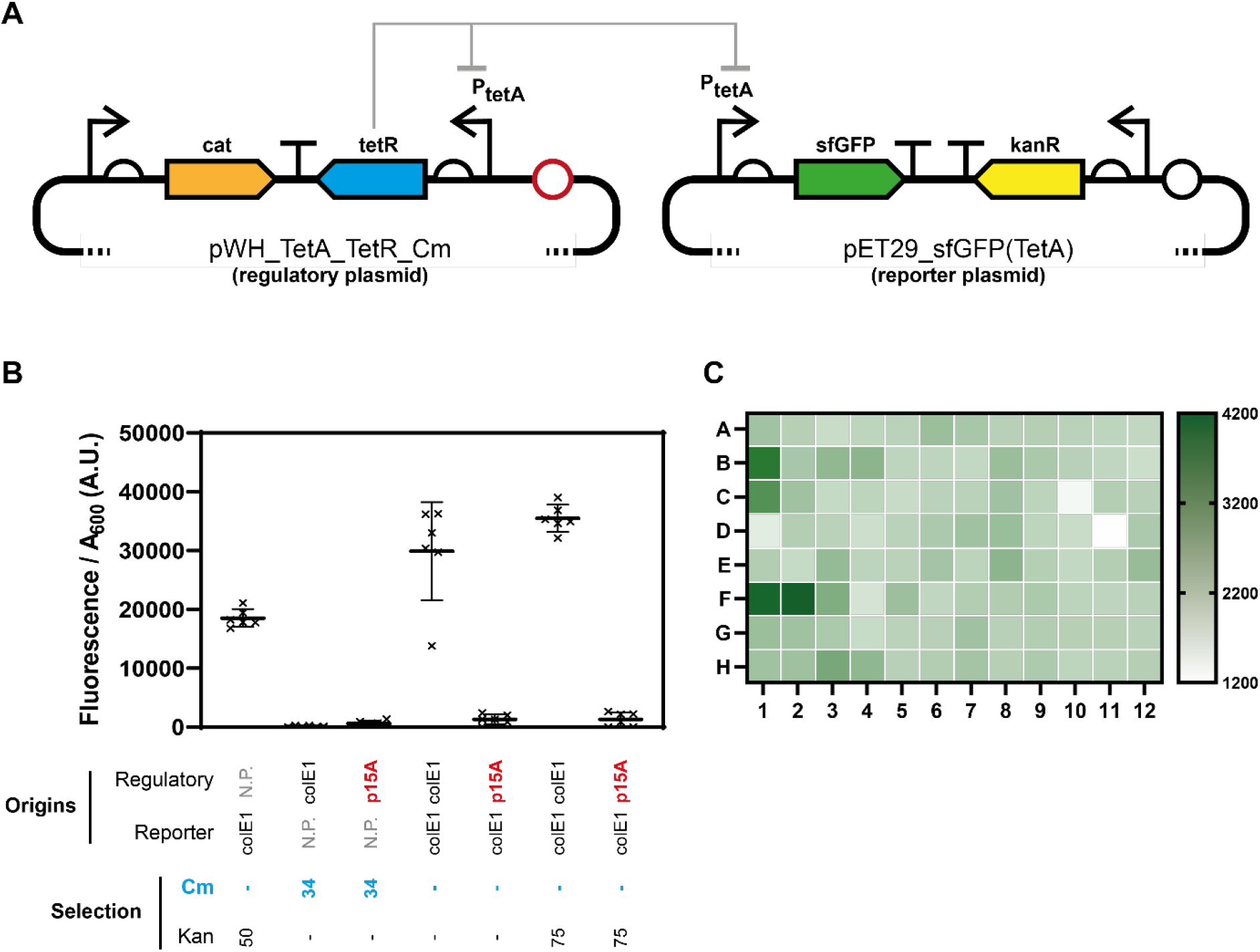
High-throughput screening of plasmid origin of replication compatibility. **A.** The 2-plasmid circuit used for the selection and screening of plasmid origin compatibility. GFP expression in the reporter plasmid (harbouring a colE1 origin – black) is inhibited by TetR being expressed from the regulatory plasmid (harbouring the tested origin – red). A negative feedback circuit (TetR expression under regulation from its own promoter) was introduced to minimise metabolic burden. Loss of regulatory plasmid deregulates the P_tetA_ promoter leading to rapid GFP expression. **B.** The circuit was validated by analysing E. coli fluorescence in culture using cells transformed with one or two plasmids after overnight growth in liquid culture supplemented with chloramphenicol (Cm), kanamycin (Kan) or no antibiotics. The compatible p15A origin from pBAD30 was used as control for a compatible origin. **C.** Heatmap of overnight selection for compatible origins. Ninety-six colonies harbouring both reporter (colE1 origin) and regulatory (origin from functional selection) plasmids were grown in the presence of kanamycin. Lowest fluorescence cultures were selected for further analysis.

By maintaining selection pressure on the reporter plasmid (i.e. the GFP coding plasmid), only the regulatory plasmid can be lost and GFP expression becomes a robust and easily quantifiable reporter for regulatory plasmid loss: accessible to both population- (spectrophotometer) and cell-based (flow cytometer) assays; and readily scalable for small volume cultures.

Small scale cultures in 96-well plates were used to validate the assay and to assess whether it could also be used as a selection tool. Overnight growth in the 96-well format (while maintaining selection for the wild-type colE1) was sufficient to identify incompatible origins (**Figure 3**) and the platform remained compatible with serial passaging, allowing longer selections (data not shown).

We therefore proceeded with the generation of all libraries for selection of plasmid viability (data not shown) and used the recovered plasmid DNA to start selection for compatible origins, introducing the library on the regulatory plasmid. Rather than harvesting all colonies for selection by serial passaging, we opted for screening a small panel directly through the high-throughput assay, as shown in **Figure 3** and **SI Fig 4**, isolating putative compatible plasmids after a single overnight growth.

Using GFP levels normalised by the density of the bacterial culture, eight isolates among the lowest GFP expression levels were selected for further analysis. Cells harbouring both wild-type and mutant origin plasmids were recovered (through dual antibiotic selection), variant origins were isolated (through transformation) and sequenced. In the case of D4, two variants were identified within the population and, once separated, were named D4 and D4II. Five of the nine variants (D4, D4II, F4, G4 and G6) were selected for detailed characterisation based on their lower sequence homology to the wild-type origin.

Plasmid compatibility of the novel origins was confirmed by serial passaging and selective plating, using selected variants in the reporter plasmid while maintaining the regulatory plasmid with its wild-type origin. Even after seven days of co-culture, significant fraction of the population retained both plasmids, as shown in **SI Fig 5**.

Notably, serial passaging in the absence of antibiotic selection tests both plasmid stability and plasmid compatibility. Biases in recovery from single antibiotic selections can be used to determine if compatibility is bidirectional. In our assays, because of the constant presence of selection for the reporter plasmid harbouring GFP, the assay tests compatibility in a single direction and its results do not address how stable the plasmids remain. Evidence of compatibility from the novel variants in the regulatory plasmids (in selection) and in the reporter plasmids (in the subsequent serial passaging) suggest that the new variants are compatible with wild-type and that this compatibility is bidirectional (in the experimental conditions used).

Although the isolated variants were selected for compatibility only against the wild-type origin, they showed significant loop variation between each other. Hence, we investigated their cross-compatibility to test if sequence divergence alone could be used as guide to assess origin compatibility.

### Cross- and inter-compatibility of novel colE1 origins

To explore all possible combinations, we generated a panel of strains harbouring all possible plasmid origin combinations between reporter and regulatory plasmids. Except for the double wild-type combination, which we repeatedly failed to successfully obtain (without the emergence of mutations within the experimental time frame), all other 35 variants were isolated.

Although successfully used to identify compatible plasmids, the high-throughput compatibility assay clearly provided high background and significant experimental variation (**SI Fig 6**). Testing a small range of alternative growth media confirmed that M9 minimal media provided the lowest fluorescence background while also maximising the discrimination between compatible and incompatible plasmids (**Figure 4** and **SI Fig 6**). Given the consistency of the assay across the different media, the assay also suggests that the isolated compatible origins remain so across a wide range of growth conditions. Flow cytometry analysis of the cultures also confirmed that the measured GFP signal was the result of high-level expression in individual cells rather than leaky GFP expression across the population (**Figure 4**).

**Figure 4:**
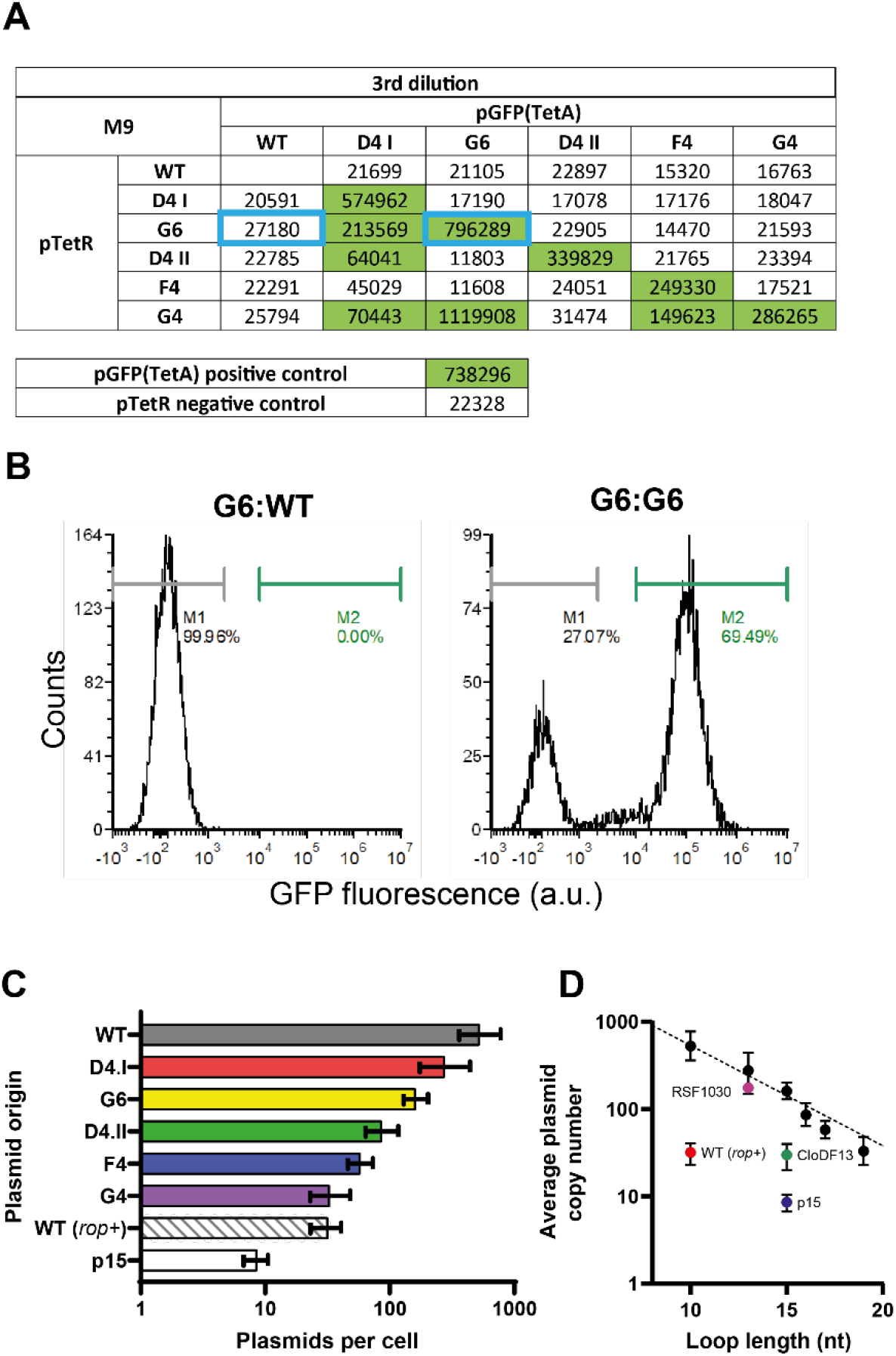
Cross-compatibility and copy number variation of colE1-compatible selected plasmid origins. **A.** Using the high-throughput assay (shown in Figure 3), the cross-compatibility between the different selected colE1-compatible plasmid origins was measured. Here, data from the third passage (therefore approximately 100 h of growth) of cells grown in M9 supplemented with kanamycin are shown. Fluorescence of each combination is shown (in arbitrary units) with significant fluorescence highlighted in green. **B.** Two origin combinations (shown as blue boxes in A) are shown here also in flow cytometry experiments, to highlight that fluorescence was the result of high GFP expression from individual cells (presumably due to regulatory plasmid loss). A more complete analysis is shown in SI Fig 6. **C.** Digital PCR results estimating the plasmid copy number of each origin in cells containing a single plasmid. Error bars represent the 95% confidence interval of the estimates. **D.** Plasmid copy numbers per cell correlates (R^2^ = 0.9818) with the sum of the RNAI loop length (based on ViennaRNA predictions), in the absence of Rom/Rop regulation.

As expected, all isolated origins remained compatible with the wild-type colE1 over multiple passages irrespective of whether they had been placed in the regulatory or reporter plasmid (**Figure 4**) – the latter maintained through antibiotic selection. The new origins also displayed the expected self-incompatibility, unrelated to copy number but presumably correlated to the dynamics of the loop interactions *in vivo*. We hypothesise that loop interactions that are not further stabilized by the hairpin rigid conformation (i.e. are not true kissing loops) lead to longer persistence of incompatible plasmids in a cell.

Cross-compatibility between the engineered origins was observed but it was not universal, with a clear dependence on which plasmid remained under selection in the assay (**Figure 4** and **SI Fig 6**). That cannot be explained by plasmid instability since that should have resulted in the systematic loss of a given origin when placed in the regulatory plasmid (not under selection).

On the other hand, given that plasmid segregation of colE1 origins is expected to be stochastic, such “directional” compatibility can be the result of differences in plasmid copy number. We therefore determined the copy number of each of the characterised origins (**Figure 4** and **SI Fig 7**). While there is correlation between plasmid loss and plasmid copy number, since plasmid loss was only observed in combinations where the regulatory plasmid was expected to be at a lower copy number than the reporter, the correlation is not perfect.

The pairwise cross-compatibility of origins also suggested plasmid combinations that should result in intercompatible origins, e.g. wild-type, D4II and G4. We tested three origin combinations using a 3-plasmid system – each expressing a different fluorescent protein (mApple, sfGFP and BFP) under an inducible promoter (**Figure 5**). Despite optimization, expression levels of blue fluorescent protein were low and discrimination between bacterial populations (expressing or not BFP) not always easy. As a result, where possible, we tracked plasmids encoding for BFP by fluorescence (experiments carried out in the absence of antibiotics) or by its resistance marker (experiments carried out in the presence of chloramphenicol).

**Figure 5:**
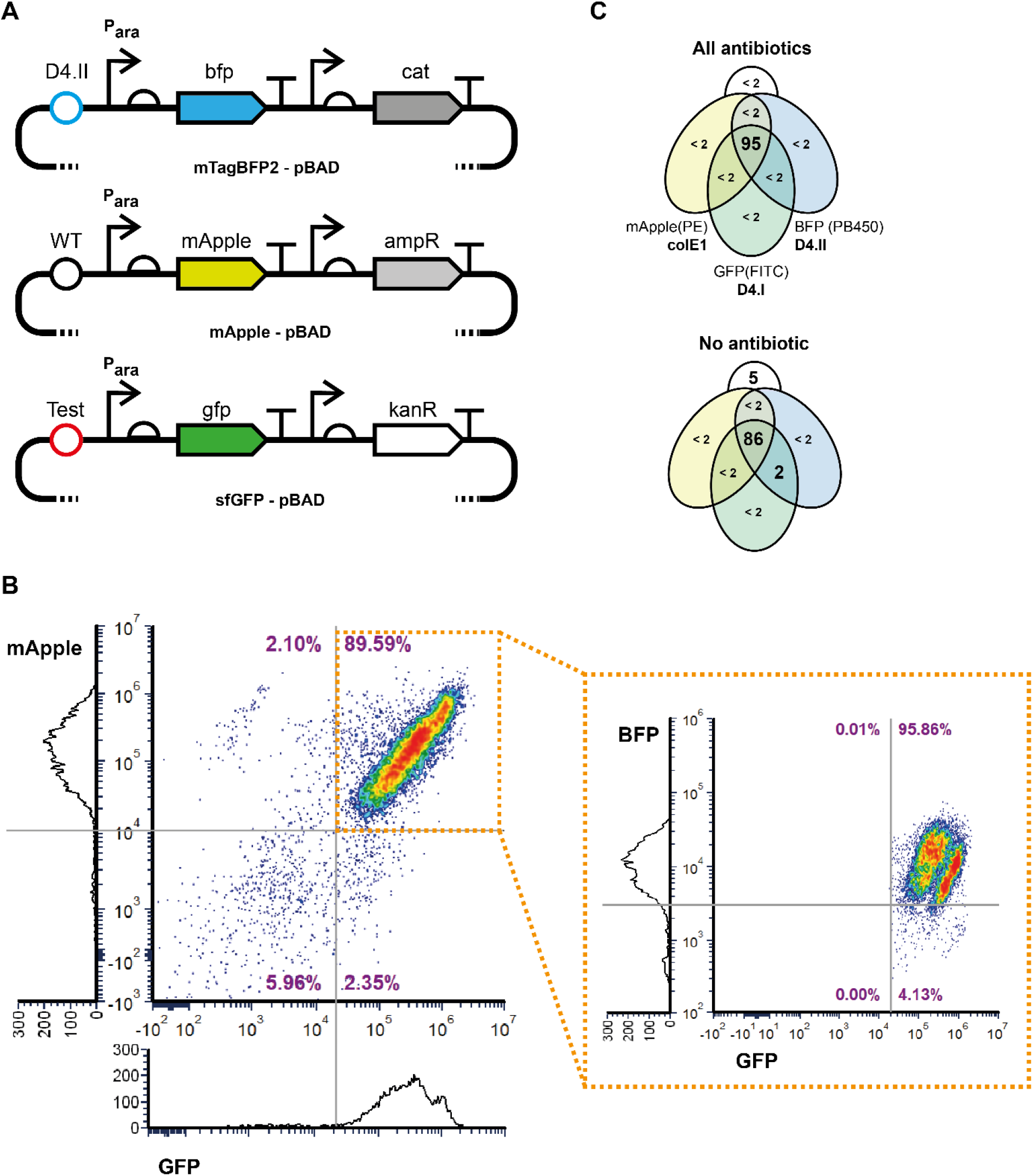
Selected plasmid origins intercompatibility. **A.** pBAD vectors harbouring fluorescent protein genes (based on plasmids # 54519, #54536 and #54572 from Addgene) were engineered to harbour different replication origins and different selectable markers. **B.** Flow cytometry analysis of bacteria harbouring colE1, D4.I and D4.II origins after overnight growth in the absence of any antibiotic selection. Cells expressing both GFP and mApple are gated (orange box) and checked for BFP expression (inset dot plot). Histograms of individual channels (without gating) are shown. **C.** Flow cytometry results are summarized in the Venn diagrams, showing all populations above 2% of the population. For the colE1, D4.I and D4.II combination, even in the absence of antibiotics, all plasmids remained stably in the cells in contrast to the two-plasmid cross-compatibility assays.

Based on the cross-compatibility assays shown in **Figure 4**, we selected two combinations we expected to be compatible (colE1/D4.I/D4.II and colE1/G6/D4.II) and one that should remain compatible under partial selection (colE1/D4.I/D4.II) for characterising their intercompatibility. While plasmid loss was, to some degree, expected in all combinations after four days of serial cultures in M9 medium, the extent of the plasmid loss differed greatly between the combinations. D4.I showed the smallest plasmid loss (7% in the absence of any antibiotics) and G4 origin the highest (40% even when all antibiotics were present) – see **SI Fig. 8** and **SI Fig. 9**. In most circumstances, plasmids with lower copy number per cell were lost, as it would be expected from a plasmid stochastic segregation process. One notable exception was seen between D4.I and wild-type (when chloramphenicol selection was maintained – **SI Fig. 9**) – where wild-type is preferentially lost despite a higher copy number – a result of more complex interactions in the regulation of the copy number of the wild-type colE1 plasmid, such as interaction with uncharged tRNAs(48). Still, the experiments confirm that pairwise cross-complementarity is not an accurate measure for higher order inter-complementarity between different origins of replication.

## Discussion and Conclusions

Despite good understanding of the colE1 mechanism of replication, there have been rare attempts at identifying and engineering colE1 plasmid compatibility, with preference frequently given to the identification of naturally occurring origins that can be tested for compatibility against colE1. Such naturally occurring origins benefit from their evolutionary-scale selection in the environment and tend to be highly stable, even in the absence of external selection, through symbiotic domestication that minimizes the metabolic cost of plasmid maintenance(49, 50). Nonetheless, it is possible to engineer compatible origins.

Our results corroborate previous findings that the loops are an important region to enable plasmid compatibility and we show that differences in loop sequence are not sufficient to predict compatibility. Plasmid maintenance assays in the absence of or under partial selection (as we show here) are an efficient route towards isolating colE1-compatible origins that remain stable in different metabolic conditions and across a wide-range of copy number.

Analysis of the predicted RNAI origins suggests a potentially novel route towards regulation of plasmid copy number, in which RNAI loop length show an inverse correlation with the estimated plasmid copy numbers per cell (**Figure 4**). This strategy, if further validated, is independent of other colE1 plasmid-based mechanisms (e.g. *cer* and *rop*), since those were not present in the plasmids used here.

Moreover, we show that pairwise compatibility assays are not sufficient to predict higher order plasmid compatibility. While the latter is hypothesised to be a consequence of the differences in plasmid copy number, copy number is not sufficient to explain the pairwise compatibility results between origins that had not been co-selected for compatibility.

Plasmid compatibility has many parallels to biological orthogonality discussed in synthetic biology(51, 52), where two biological systems need to co-exist *in vivo* without significant interaction. We propose, in fact, that it can be used as a model to improve the concept of orthogonality, since plasmid orthogonality is nuanced: operating on a scale rather than in absolutes, affected by experimental conditions and having to operate within the more complex cellular milieu.

Although the current focus of biomanufacturing still lies on improving plasmid stability, programmable plasmid compatibility can potentially lead to novel applications or to new strategies of biological containment. Our work is a first step towards bespoke plasmid origins and programmable plasmid orthogonality.

## Acknowledgements

VBP, LT and WH thank ERASynBio (grant BB/N01023X/1; *invivo*XNA). VBP and ABR thank ERASynBio (grant BB/N010221/1; TNAepisome). VBP and SC thank the Rega Foundation and KU Leuven (grant C14/19/102).

## Contributions

VBP, ES and SC contributed to the project design. ES performed all early large-scale experiments, initial selections for viability. ES and VBP carried out the NGS analysis. VBP designed, WH and SC assembled the required biological circuits, and ABR developed and validated the high-throughput plasmid compatibility assay. LT and SC carried out selections for plasmid compatibility. SC characterised isolated origins and investigated cross- and inter-compatibility between isolated origins. SC and VBP wrote the manuscript.

## Bibliography

1. Staal, J., Alci, K., De Schamphelaire, W., Vanhoucke, M. and Beyaert, R. (2019) Engineering a minimal cloning vector from a pUC18 plasmid backbone with an extended multiple cloning site. Biotechniques, 66, 254–259.

2. Johnston, C.D., Cotton, S.L., Rittling, S.R., Starr, J.R., Borisy, G.G., Dewhirst, F.E. and Lemon, K.P. (2019) Systematic evasion of the restriction-modification barrier in bacteria. Proc. Natl. Acad. Sci., 116, 11454–11459.

3. Martínez-García, E., Goñi-Moreno, A., Bartley, B., McLaughlin, J., Sánchez-Sampedro, L., Pascual Del Pozo, H., Prieto Hernández, C., Marletta, A.S., De Lucrezia, D., Sánchez-Fernández, G., et al. (2020) SEVA 3.0: An update of the Standard European Vector Architecture for enabling portability of genetic constructs among diverse bacterial hosts. Nucleic Acids Res., 48, D1164–D1170.

4. Silva-Rocha, R., Martínez-García, E., Calles, B., Chavarría, M., Arce-Rodríguez, A., De Las Heras, A., Páez-Espino, A.D., Durante-Rodríguez, G., Kim, J., Nikel, P.I., et al. (2013) The Standard European Vector Architecture (SEVA): A coherent platform for the analysis and deployment of complex prokaryotic phenotypes. Nucleic Acids Res., 41, D666.

5. Valenzuela-Ortega, M. and French, C. (2021) Joint universal modular plasmids (JUMP): a flexible vector platform for synthetic biology. Synth. Biol., 6.

6. Hasty, J., McMillen, D. and Collins, J.J. (2002) Engineered gene circuits. Nature, 420, 224–230.

7. Morgan, J.L.W., Acheson, J.F. and Zimmer, J. (2017) Structure of a Type-1 Secretion System ABC Transporter. Structure, 25, 522–529.

8. Romier, C., Ben Jelloul, M., Albeck, S., Buchwald, G., Busso, D., Celie, P.H.N., Christodoulou, E., De Marco, V., van Gerwen, S., Knipscheer, P., et al. (2006) Co-expression of protein complexes in prokaryotic and eukaryotic hosts: experimental procedures, database tracking and case studies. Acta Crystallogr. Sect. D Biol. Crystallogr., 62, 1232–1242.

9. Yang, J. and Guo, L. (2014) Biosynthesis of β-carotene in engineered E. coli using the MEP and MVA pathways. Microb. Cell Fact., 13, 1–11.

10. Martin, V.J.J., Piteral, D.J., Withers, S.T., Newman, J.D. and Keasling, J.D. (2003) Engineering a mevalonate pathway in Escherichia coli for production of terpenoids. Nat. Biotechnol., 21, 796–802.

11. Wang, L. and Schultz, P.G. (2001) A general approach for the generation of orthogonal tRNAs. Chem. Biol., 8, 883–890.

12. Packer, M.S., Rees, H.A. and Liu, D.R. (2017) Phage-assisted continuous evolution of proteases with altered substrate specificity. Nat. Commun., 8, 956.

13. Velappan, N., Sblattero, D., Chasteen, L., Pavlik, P. and Bradbury, A.R.M. (2007) Plasmid incompatibility: more compatible than previously thought? Protein Eng. Des. Sel., 20, 309–313.

14. Novick, R.P. (1987) Plasmid incompatibility. Microbiol. Rev., 51, 381–395.

15. Shintani, M., Sanchez, Z.K. and Kimbara, K. (2015) Genomics of microbial plasmids: Classification and identification based on replication and transfer systems and host taxonomy. Front. Microbiol., 6, 242.

16. Selzer, G., Som, T., Itoh, T. and Tomizawa, J. (1983) The origin of replication of plasmid p15A and comparative studies on the nucleotide sequences around the origin of related plasmids. Cell, 32, 119–129.

17. Morita, M. and Oka, A. (1979) The structure of a transcriptional unit on colicin E1 plasmid. Eur. J. Biochem., 97, 435–43.

18. Lacatena, R.M. and Cesareni, G. (1981) Base pairing of RNA I with its complementary sequence in the primer precursor inhibits ColE1 replication. Nature, 294, 623–626.

19. Cesareni, G., Helmer-Citterich, M. and Castagnoli, L. (1991) Control of ColE1 plasmid replication by antisense RNA. Trends Genet., 7, 230–235.

20. Cesareni, G., Muesing, M.A. and Polisky, B. (1982) Control of ColE1 DNA replication: the rop gene product negatively affects transcription from the replication primer promoter. Proc. Natl. Acad. Sci., 79, 6313–6317.

21. Salim, N., Lamichhane, R., Zhao, R., Banerjee, T., Philip, J., Rueda, D. and Feig, A.L. (2012) Thermodynamic and kinetic analysis of an RNA kissing interaction and its resolution into an extended duplex. Biophys. J., 102, 1097–1107.

22. Kim, D., Rhee, Y., Rhodes, D., Sharma, V., Sorenson, O., Greener, A. and Smider, V. (2005) Directed Evolution and Identification of Control Regions of ColE1 Plasmid Replication Origins Using Only Nucleotide Deletions. J. Mol. Biol., 351, 763–775.

23. Tomizawa, J. and Itoh, T. (1981) Plasmid ColE1 incompatibility determined by interaction of RNA I with primer transcript. Proc. Natl. Acad. Sci., 78, 6096–6100.

24. Camps, M. (2010) Modulation of ColE1-like plasmid replication for recombinant gene expression. Recent Patents DNA Gene Seq., 4, 58–73.

25. de Lorenzo, V. (2011) Beware of metaphors: Chasses and orthogonality in synthetic biology. Bioeng. Bugs, 2, 3–7.

26. Liu, C.C., Jewett, M.C., Chin, J.W. and Voigt, C.A. (2018) Toward an orthogonal central dogma. Nat. Chem. Biol., 14, 103–106.

27. Costello, A. and Badran, A.H. (2021) Synthetic Biological Circuits within an Orthogonal Central Dogma. Trends Biotechnol., 39, 59–71.

28. Torres, L., Krüger, A., Csibra, E., Gianni, E. and Pinheiro, V.B.B. (2016) Synthetic biology approaches to biological containment: Pre-emptively tackling potential risks. Essays Biochem., 60.

29. Neumann, H., Slusarczyk, A.L. and Chin, J.W. (2010) De novo generation of mutually orthogonal aminoacyl-tRNA synthetase/tRNA pairs. J. Am. Chem. Soc., 132, 2142–4.

30. McClune, C.J., Alvarez-Buylla, A., Voigt, C.A. and Laub, M.T. (2019) Engineering orthogonal signalling pathways reveals the sparse occupancy of sequence space. Nature, 574, 702–706.

31. Padgett, K.A. and Sorge, J.A. (1996) Creating seamless junctions independent of restriction sites in PCR cloning. Gene, 168, 31–35.

32. Gibson, D.G., Glass, J.I., Lartigue, C., Noskov, V.N., Chuang, R.-Y., Algire, M.A., Benders, G.A., Montague, M.G., Ma, L., Moodie, M.M., et al. (2010) Creation of a Bacterial Cell Controlled by a Chemically Synthesized Genome. Science (80-.)., 329, 52–56.

33. Chen, W. and Kuo, T. (1993) A simple and rapid method for the preparation of gram-negative bacterial genomic DNA. Nucleic Acids Res., 21, 2260–2260.

34. Bhat, S., Herrmann, J., Armishaw, P., Corbisier, P. and Emslie, K.R. (2009) Single molecule detection in nanofluidic digital array enables accurate measurement of DNA copy number. Anal. Bioanal. Chem., 394, 457–467.

35. Heid, C.A., Stevens, J., Livak, K.J. and Williams, P.M. (1996) Real time quantitative PCR. Genome Res., 6, 986–994.

36. Afgan, E., Baker, D., van den Beek, M., Blankenberg, D., Bouvier, D., Čech, M., Chilton, J., Clements, D., Coraor, N., Eberhard, C., et al. (2016) The Galaxy platform for accessible, reproducible and collaborative biomedical analyses: 2016 update. Nucleic Acids Res., 44, W3–W10.

37. Lorenz, R., Bernhart, S.H., Höner zu Siederdissen, C., Tafer, H., Flamm, C., Stadler, P.F. and Hofacker, I.L. (2011) ViennaRNA Package 2.0. Algorithms Mol. Biol., 6, 26.

38. Gruber, A.R., Lorenz, R., Bernhart, S.H., Neubock, R. and Hofacker, I.L. (2008) The Vienna RNA Websuite. Nucleic Acids Res., 36, W70–W74.

39. Kerpedjiev, P., Hammer, S. and Hofacker, I.L. (2015) Forna (force-directed RNA): Simple and effective online RNA secondary structure diagrams. Bioinformatics, 31, 3377–3379.

40. Tizei, P.A.G., Csibra, E., Torres, L. and Pinheiro, V.B. (2016) Selection platforms for directed evolution in synthetic biology. Biochem. Soc. Trans., 44.

41. Masukata, H. and Tomizawa, J. (1990) A mechanism of formation of a persistent hybrid between elongating RNA and template DNA. Cell, 62, 331–338.

42. Lorenz, R., Bernhart, S.H., Höner zu Siederdissen, C., Tafer, H., Flamm, C., Stadler, P.F. and Hofacker, I.L. (2011) ViennaRNA Package 2.0. Algorithms Mol. Biol., 6, 1–14.

43. Hoesl, M.G., Oehm, S., Durkin, P., Darmon, E., Peil, L., Aerni, H.-R., Rappsilber, J., Rinehart, J., Leach, D., Söll, D., et al. (2015) Chemical Evolution of a Bacterial Proteome. Angew. Chemie Int. Ed., 54, 10030–10034.

44. Bao, L., Menon, P.N.K., Liljeruhm, J. and Forster, A.C. (2020) Overcoming chromoprotein limitations by engineering a red fluorescent protein. Anal. Biochem., 611, 113936.

45. Shaner, N.C., Steinbach, P.A. and Tsien, R.Y. (2005) A guide to choosing fluorescent proteins. Nat. Methods, 2, 905–909.

46. Jahn, M., Vorpahl, C., Hübschmann, T., Harms, H. and Müller, S. (2016) Copy number variability of expression plasmids determined by cell sorting and Droplet Digital PCR. Microb. Cell Fact., 15, 211.

47. Grabherr, R., Nilsson, E., Striedner, G. and Bayer, K. (2002) Stabilizing plasmid copy number to improve recombinant protein production. Biotechnol. Bioeng., 77, 142–147.

48. Wróbel, B. and Wę, G. (1998) Replication Regulation of ColE1-like Plasmids in Amino Acid-StarvedEscherichia coli. Plasmid, 39, 48–62.

49. Wein, T., Hülter, N.F., Mizrahi, I. and Dagan, T. (2019) Emergence of plasmid stability under non-selective conditions maintains antibiotic resistance. Nat. Commun., 10, 2595.

50. MacLean, R.C. and San Millan, A. (2015) Microbial Evolution: Towards Resolving the Plasmid Paradox. Curr. Biol., 25, R764–R767.

51. Liu, C.C., Jewett, M.C., Chin, J.W. and Voigt, C.A. (2018) Toward an orthogonal central dogma. Nat. Chem. Biol., 14, 103–106.

52. de Lorenzo, V. (2011) Beware of metaphors: Chasses and orthogonality in synthetic biology. Bioeng. Bugs, 2, 3–7.

