## Supplementary information for "Directed evolution of colE1 plasmid replication compatibility: a fast tractable tunable model for investigating biological orthogonality"

* contributed equally to the publication.

**Supplementary information:**

Supplementary Table 1: Primers used in this work ... 3

Supplementary Table 2: Number of events post single-cell gating used in the analysis of plasmid populations. ... 3

Supplementary Figure 1: Viable colE1 origins identified by NGS and screening. ... 4

Supplementary Table 3: Analysis by next generation sequencing of recovered viable origins. ... 4

Supplementary Figure 2: Compatibility selection in liquid culture. ... 5

Supplementary Figure 3: NGS analysis after large-scale selection for plasmid compatibility. ... 5

Supplementary Table 4: Analysis by next generation sequencing of recovered compatible origins. ... 6

Supplementary Figure 4: High-throughput screening assay for the selection of colE1-compatible origins of replication. ... 6

Supplementary Figure 5: Characterisation of selected colE1 origin variants for their compatibility with colE1. ... 7

Supplementary Figure 6: Impact of culture medium on plasmid compatibility and characterization of cross-compatibility. ... 8

Supplementary Figure 7: Sequence analysis of engineered colE1 origins and quantification of plasmid copy number per cell. ... 9

Supplementary Figure 8: Pairwise compatibility between origins in plasmids expressing fluorescent proteins. ... 10

Supplementary Figure 9: Plasmid intercompatibility assays. ... 11

Supplementary notes: ... 12

| Primer name | Primer sequence |
|---|---|
| **Construct and library assembly** | |
| ES VCTR FW | AAAGGTCTCAAGTGTAGCCGTAGTTAGGC |
| ES VCTR RV | AAAGGTCTCACAAGCAGCAGATTACGCG |
| ES ULTR RNAI FW | GGGGGTCTCACACTTAGAAG |
| ES ULTR REV | ACACACAGGTCTCACTTGC |
| ES COLEI ins GIBS FW | CTACGCATGGCTCAAAACACCCCTTGT |
| ES COLEI ins GIBS RV | TTTTTCCATAGGCTCCGCC |
| ES 002 FW | AAAGGTCTCACGTTAAAGGAAGCTGAGTTGGCT |
| ES 002 RV | AAAGGTCTCATAGAGGGGAATTGTTATCCGC |
| ES 003 FW | AAAGGTCTCATCTAGGGCTAACAGGAGGAATTAAC |
| ES 003 RV | AAAGGTCTCAAACGCATCCGCCAAAACAGC |
| ES BAD GFP FW | AAAGGTCTCACCCGTTTTTTGGGCTAAC |
| ES BAD GFP RV | AAAGGTCTCAGCTTCGCTTCTGCGTTCTGAT |
| ES PET GFP FW | AAAGGTCTCAAAGCCCGAAAGGAA |
| ES PET GFP RV | AAAGGTCTCACGGGAATTGTTATCCGCT |
| ES sfGFP plasm ampl FW | GGCGGAGCCTATGGAAAAA |
| ES sfGFP plasm ampl RV | GGGGTGTTTTGAGCCATGCGTAGAGGATCTGCTCA |
| SC_pBAD_dOri_FW | AAACGTCTCACTTGCATGTGTCAGAGGTTTTCAC |
| SC_pBAD_dOri_RV | AAACGTCTCATCACTCAGTGGAACGAAAACTCAC |
| SC_pBAD_dOri_dATB_RV | AAACGTCTCATCACTGTAGAAACGCAAAAAGGCC |
| SC_pET_addOri_addATB_FW | AAAGGTCTCAGTGACGTTTACAATTTCAGGTGGC |
| SC_pET_addOri_addATB_RV | AAAGGTCTCACAAGATCAGCTCACTCAAAGGC |
| SC_pWH_addOri_addATB_FW | AAAGGTCTCAGTGATTCCGTGATGGTAACTTCAC |
| SC_pWH_addOri_FW | AAAGGTCTCAGTGAGCAAGGATCTTCTTGAGATCC |
| SC_pWH_addOri_addATB_RV | AAAGGTCTCACAAGAATCATCTGGCCATTCGATG |
| AR_pWHalpha_Fw | TATGGAAAAACGCCAGCAACG |
| AR_pWHalpha_Rv | AAGATCCTTGCACTCGAGTTGATCG |
| VP023F | TTTGGTCTCA AAGTTGCACTCGAGTTGATCGGGC |
| VP023R | TTTGGTCTCA TTCCGCCTTTTTACGGTTCCTGGCC |
| WH81 | TCCTCGAGGCTTGGATTCTC |
| WH82 | TGCACTCGAGTTGATCGGG |
| WH83 | TGCCCGATCAACTCGAGTGCAAGGATCTTCTTGAGATCC |
| WH84 | AACGAGACATCATTTTTTGCCCTCGTTATCTAG |
| WH85 | AGGGCAAAAAATGATGTCTCGTTTAGATAAAAG |
| WH86 | AGAATCCAAGCCTCGAGGAAGATCCTTTGATCTTTTCTAC |
| WH87 | TCCCTATCAGTGATAGAGAACCTCTAGAAATAATTTTGTTTAAC |
| WH88 | ATCAATGATAGAGTGTCAACATTTCGCGGGATCGAG |
| TetA | GTTGACACTCTATCATTGATAGAGTTATTTTACCACTCCCTATCAGTGATAGAGAA |
| TetAR | TTCTCTATCACTGATAGGGAGTGGTAAAATAACTCTATCAATGATAGAGTGTCAAC |
| Sequencing | |
| ES seq-ing 001 | TCACTCAAAGGCGGTAA |
| ES seq-ing 002 | TGTCGGGTCATGTGAGCAA |
| ES seq-ing 002 FW | ATGGCTCATAACACCCCTTGT |
| NGS_Forward primer | TTCTGCGCGTAATCTGCTGC |
| NGS_Reverse primer | GGCCTAACTACGGCTACACTAG |
| ES DEEP SEQ PET ini FW | AATGATACGGCGACCACCGAGATCTACACTCTTTCCCTACACGACGCTCTTCCGATCTNNNT GACCATTCTGCGCGTAATCTGCTGC |
| ES DEEP SEQ PET vai FW | AATGATACGGCGACCACCGAGATCTACACTCTTTCCCTACACGACGCTCTTCCGATCTNNN ACAGTGTTCTGCGCGTAATCTGCTGC |
| ES DEEP SEQ PET RV | CAAGCAGAAGACGGCATACGAGATCGGTCTCGGCATTCCTGCTGAACCGCTCTTCCGATCTG GCCTAACTACGGCTACACTAG |
| Digital PCR | |

| | |
|---|---|
| SC_dPCR_Chl_FW | AATAAAGGCCGGATAAAACTTG |
| SC_dPCR_Chl_RV | CTGGATATACCACCGTTGATAT |
| SC_dPCR_Chl_probe | /56-FAM/AATATCCAG/ZEN/CTGAACGGTCTGG/3IABKFQ/ |
| SC_dPCR_ter_FW | AATAACATTCATTGGGTTGGTC |
| SC_dPCR_ter_RV | GCATGGTTAATCACGATGTAAT |
| SC_dPCR_ter_probe | /5HEX/AATAGCTAC/ZEN/CTCATCCGCGAAG/3IABKFQ/ |

**Supplementary Table 1: Primers used in this work.** All primers are shown in 5'→3' orientation. Chemical modifications for the primers used in digital PCR were as follows: /56-FAM/ - fluorescein; /ZEN/ - ZEN™ quencher; /3IABKFQ/ - Iowa Black® FQ; /5HEX/ - Hexachlorofluorescein.

| Plasmid combination | # events |
|---|---|
| **Intercompatibility experiments** | |
| **D4_1 (all)** | 2754 |
| **D4_1 (CM only)** | 43973 |
| **D4_1 (no ATB)** | 9029 |
| **G6 (all)** | 6736 |
| **G6 (CM only)** | 10225 |
| **G6 (no ATB)** | 3597 |
| **G4 (all)** | 2417 |
| **G4 (CM only)** | 9626 |
| **G4 (no ATB)** | 3046 |
| **Pairwise_intercompatibility experiments** | |
| **D4_2 + colE1 (CM)** | 8804 |
| **D4_2 + colE1 (no ATB)** | 6896 |
| **D4_1 + colE1 (no ATB)** | 1520 |
| **colE1 + G4 (no ATB)** | 8195 |
| **colE1 + G6 (no ATB)** | 5304 |
| **D4_1 + D4_2 (CM only)** | 1903 |
| **D4_1 + D4_2 (no ATB)** | 6851 |
| **D4_2 + G6 (CM only)** | 2625 |
| **D4_2 + G6 (no ATB)** | 9647 |
| **D4_2 + G4 (CM only)** | 1399 |
| **D4_2 + G4 (no ATB)** | 2428 |

**Supplementary Table 2: Number of events post single-cell gating used in the analysis of plasmid populations.** Naming of the experiments refers to the origins from each plasmid as described in the main text. Compatibility experiments carried out in the presence of kanamycin, chloramphenicol and ampicillin are shown as (all). Where only chloramphenicol was used in the experiment, samples are shown as (CM only). Experiments carried out in the absence of any antibiotic are shown as (no ATB).

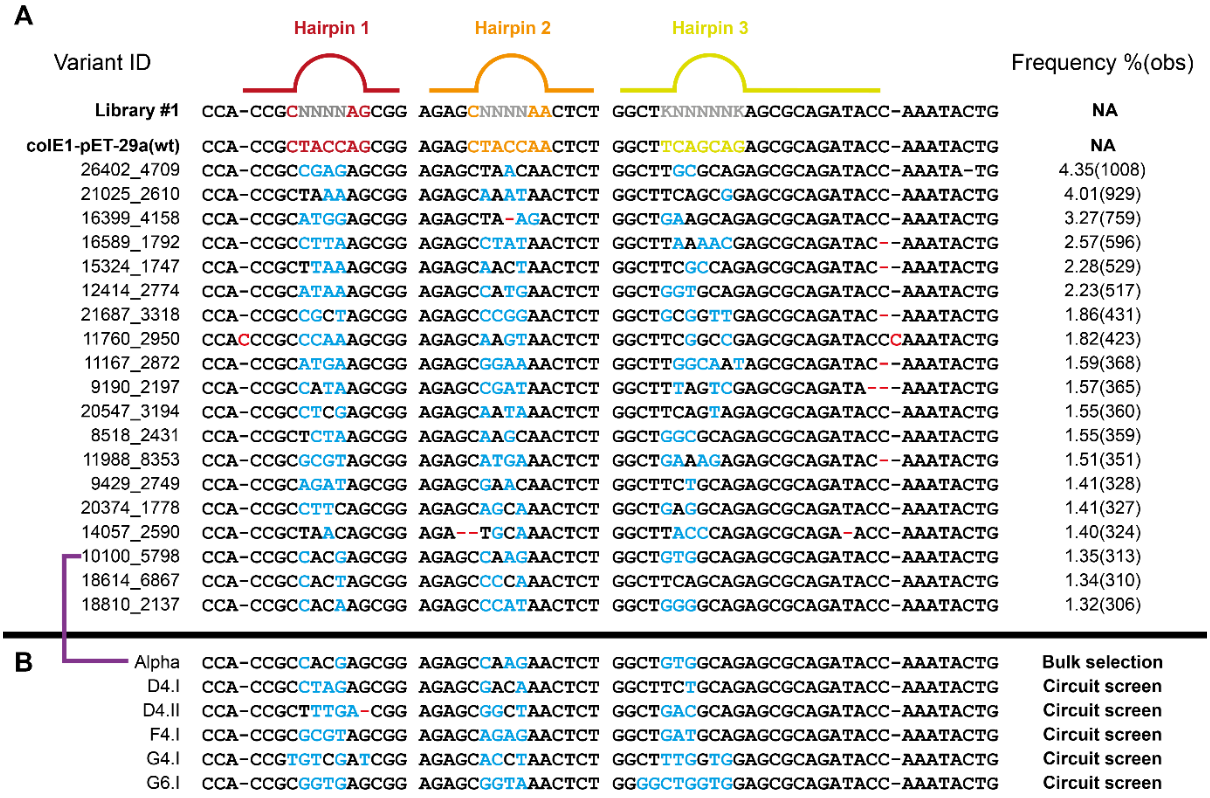

**Supplementary Figure 1: Viable colE1 origins identified by NGS and screening. A.** NGS analysis of viable colE1 origins isolated from transformation of library #1. Mutations away from the wild-type sequence introduced by the library are shown in blue, mutations arising from selection are shown in red. Frequency of isolated origins is shown with the individual number of observations in brackets. The ID (automatically generated in sequencing) of one of the unique sequences is picked (arbitrarily) to name the group. NA – not applicable. **B.** Engineered colE1 origins described in this work.

| Pipeline step | Sequences output |
|---|---|
| Total read number | 31144 |
| Quality trimming | 31144 (100%) |
| Filtering by 5' sequence | 29916 (96%) |
| Filtering by 5' sequence #2 | 29151 (94%) |
| Filtering by 3' sequence | 27175 (87%) |
| Filtering by 3' sequence #2 | 23183* (74%) |
| Unique sequences | 1903 |

**Supplementary Table 3: Analysis by next generation sequencing of recovered viable origins.** Total read number obtained and the impact of the analysis pipeline are shown. *Number of sequences used in downstream analysis.

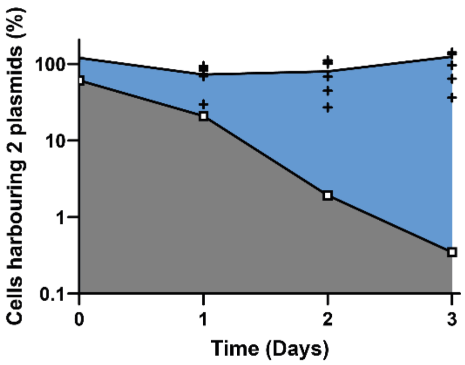

**Supplementary Figure 2: Compatibility selection in liquid culture.** *E. coli* harbouring pSB1C3 (colE1 origin) and transformed with pET29 containing its wild-type (colE1; white squares) or a library of viable origins (black crosses) were serially passaged, with samples plated in the absence of antibiotics (to determine total CFU) or in the presence of both antibiotics (to determine CFU still harbouring both plasmids). As expected, under the growth conditions, the wild type colE1 origin is rapidly lost from the population.

**A**

| Variant ID | Hairpin 1 | Hairpin 2 | Hairpin 3 | Frequency %(obs) |
|---|---|---|---|---|
| **colE1_pET-29a(wt)** | CCACCGCTACCAGCGG | AGAGCTACCAACTCT | GGCTTCAGCAGAGCGCAGATACCAAATACTG | **NA** |
| 19921_1932 | CCACCGCTACCAGCGG | AGAGCTACCAACTCT | GGCTTCAGCAGAGCGCAGATACCAAATACTG | (13432) |
| 24147_3005 (alpha) | CCACCGCCACGAGCGG | AGAGCCAAGAACTCT | GGCTGTGGCAGAGCGCAGATACCAAATACTG | 11.22(1433) |
| 16596_2335 | CCACCGCACCGAGCGG | AGAGCCGCCAACTCT | GGTTGACTAAGAGCGCAGATACCAAATACTG | 4.01(512) |
| 15490_2347 | CCACCGCCATAAGCGG | AGAGCCGCCAACTCT | GGCTGGCAGAGAGCGCAGATACCAAATACTG | 3.63(464) |
| 23482_1919 | CCACCGCAGAGAGCGG | AGAGCCCAAAACTCT | GGCTTCGGCAGAGCGCAGATACCAAATACTG | 3.58(457) |
| 11945_7277 | CCACCGCACCTAGCGG | AGAGCCATCAACTCT | GGCTGACGCAGAGCGCAGATACCAAATACTG | 3.52(450) |
| 21796_3950 | CCACCGCACATAGCGG | AGAGCTTTCAACTCT | GGCTTGAGCAGAGCGCAGATACCAAATACTG | 3.21(410) |
| 6435_3326 | CCACCGCCACTAGCGG | AGAGCCCCAAACTCT | GGCTTCAGCAGAGCGCAGATACCAAATACTG | 2.72(347) |
| 13190_7993 | CCACCGCAGAGAGCGG | AGAGCCCTCAACTCT | GGCTGTTGCAGAGCGCAGATACCAAATACTG | 2.51(321) |
| 13128_4999 | CCACCGCCCACAGCGG | AGAGCCAGCAACTCT | GGCTGAGTAAGAGCGCAGA-ACCAAATACTG | 2.47(315) |
| 13613_5267 | CCACCGCCACGAGCGG | AGAGCTACAAACTCT | GGCTTCGACAGAGCGCAGA-ACCAAATACTG | 2.43(310) |
| 16898_3487 | CCACCGCCATGAGCGG | AGAGCCCGCAACTCT | GGCTTGACCTGAGCGCAGATA-CAAATACTG | 1.97(252) |
| 25074_2902 | CCACCGCAGACAGCGG | AGAGCCTCTAACTCT | GGCTGACGCTGAGCGCAGA-ACCAAATACTG | 1.96(251) |
| 21215_6358 | CCACCGCACCAAGCGG | AGAGCGCTTAACTCT | GGCTTACGCAGAGCGCAGATACCAAATACTG | 1.96(250) |
| 12850_3238 | CCACCGCCCTAAGCGG | --AGCCATTAACTCT | GGCTTATGCAGAGCGCAGATACCAAATACTG | 1.78(227) |
| 25300_8619 | CCACCGCAGATAGCGG | AGAGCCAACAACTCT | GGCTTCCGCAGAGCGCAGATACCAAATACTG | 1.76(225) |
| 14580_2678 | CCACCGCCACCAGCGG | AGAGCTATTAACTCT | GGCTTCGGCAGAGCGCAGATACCAAATACTG | 1.73(221) |

**Supplementary Figure 3: NGS analysis after large-scale selection for plasmid compatibility.** NGS analysis of colE1 origins isolated after selection of viable colE1 variants co-transformed with wild-type colE1. Mutations away from the wild-type sequence introduced by the library are shown in blue, mutations arising from selection are shown in red. Frequency of isolated origins is shown with the individual number of observations in brackets. Wild-type colE1 sequences were identified in the experiment (a limitation of the approach used to prepare plasmid DNA for NGS) and are excluded from the analysis – the number of observations is still given. The ID (automatically generated in sequencing) of one the unique sequences is picked (arbitrarily) to name the group. NA – not applicable.

| Pipeline step | Sequences output |
| --- | --- |
| **Total read number** | 28438 |
| **Quality trimming** | 28438 (100%) |
| **Filtering by 5' sequence** | 27876 (98%) |
| **Filtering by 5' sequence #2** | 27708 (97%) |
| **Filtering by 3' sequence** | 26777 (94%) |
| **Filtering by 3' sequence #2** | 26206* (92%) |
| **Unique sequences** | 1185 |

**Supplementary Table 4: Analysis by next generation sequencing of recovered compatible origins.** Total read number obtained and the impact of the analysis pipeline are shown. *Number of sequences used in downstream analysis.

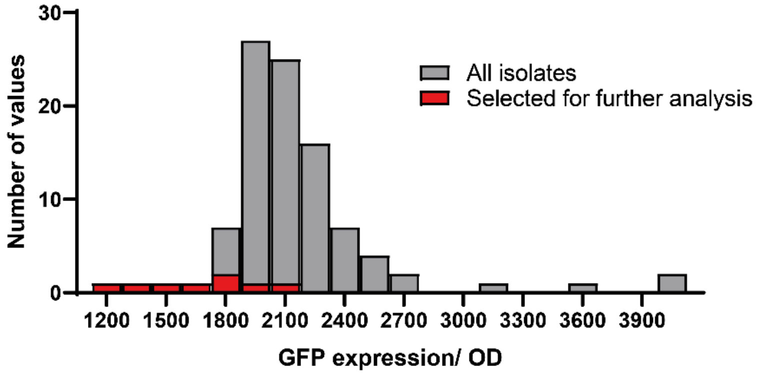

**Supplementary Figure 4: High-throughput screening assay for the selection of colE1-compatible origins of replication.** Histogram representation of the data shown in Figure 3C, highlighting the fluorescence values of the variants selected for further study.

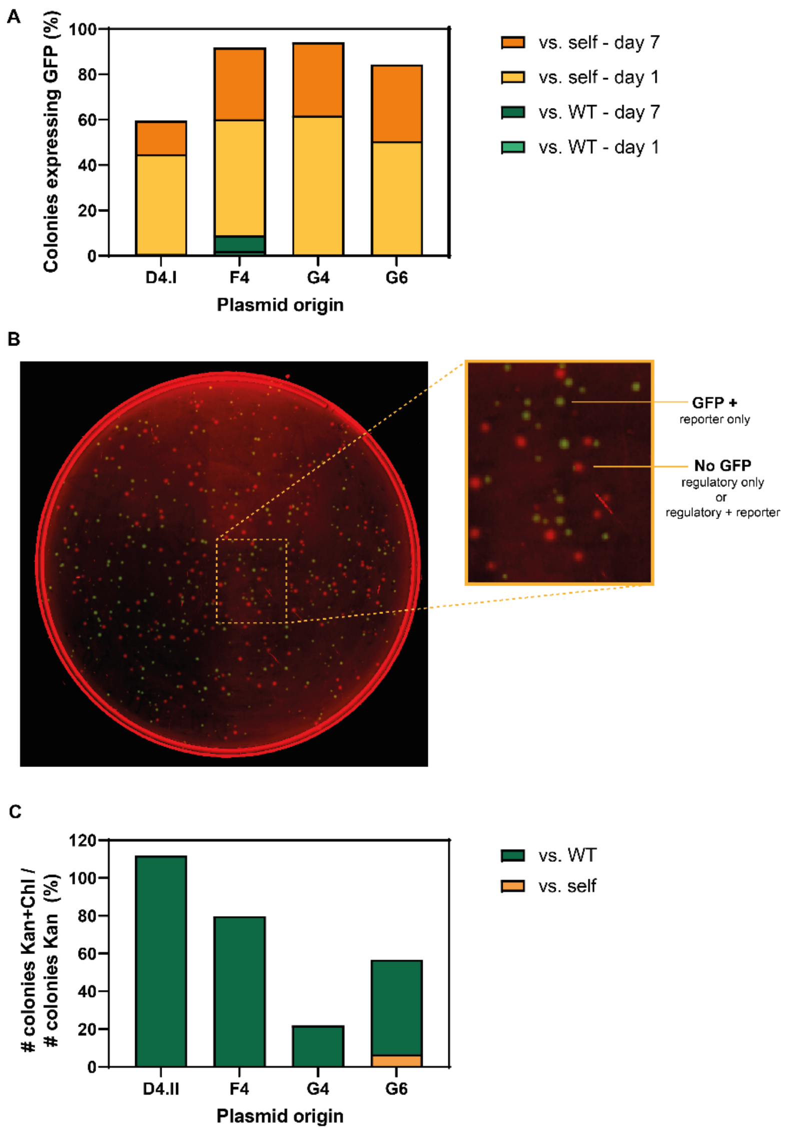

**Supplementary Figure 5: Characterisation of selected colE1 origin variants for their compatibility with colE1. A.** Serial cultures of *E. coli* cells co-transformed with reporter (harbouring one of D4.I, F4, G4, G6 and wild-type origins of replication) and regulatory (harbouring D4.I, F4, G4 or G6 origins) were used to test the compatibility of the selected variants against wild-type and to confirm their self-incompatibility. The percentage of cells expressing GFP was calculated by diluting a culture aliquot (after 1 or 7 days of passaging) and plating in LB agar supplemented with chloramphenicol (to retain reporter plasmid). Bar graphs are overlaid with compatibility to wild-type shown in green (light or dark depending on passage number) and self-incompatibility shown in orange (light or dark depending on passage number). **B.** Example of transformation plate used to calculate values in **A.** Here, D4.I was used in both regulatory and reporter plasmids and the results show the distribution of plasmids after 7 days of passaging. Fluorescent images from GFP (green) and control channels (red) are overlaid and CFU counted. **C.** Complementary experiment where after passaging in the absence of antibiotic selection, cultures are plated in media supplemented with kanamycin (regulatory plasmid antibiotic marker) or with both antibiotics to monitor plasmid loss. Bar graphs are overlaid with compatibility to wild-type shown in green and self-incompatibility shown in orange.

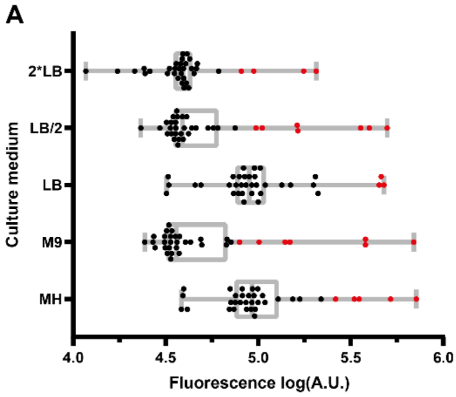

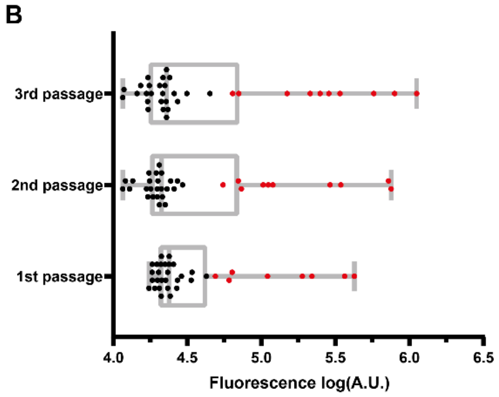

**C**

| M9 | | 1st dilution pGFP(TetA) WT | D4 I | G6 | D4 II | F4 | G4 |
|---|---|---|---|---|---|---|---|
| pTetR | WT | | 28838 | 27170 | 24209 | 23776 | 23285 |
| | D4 I | 19704 | 34078 | 25503 | 18343 | 20212 | 20976 |
| | G6 | 23319 | 220142 | 365037 | 22609 | 20490 | 22387 |
| | D4 II | 21133 | 42589 | 21106 | 60662 | 18349 | 18293 |
| | F4 | 19862 | 26849 | 21184 | 19061 | 63579 | 17417 |
| | G4 | 33635 | 110283 | 425005 | 23830 | 48928 | 189417 |

| | |
|---|---|
| pGFP(TetA) positive control | 387180 |
| pTetR negative control | 19953 |

| M9 | | 3rd dilution pGFP(TetA) WT | D4 I | G6 | D4 II | F4 | G4 |
|---|---|---|---|---|---|---|---|
| pTetR | WT | | 21699 | 21105 | 22897 | 15320 | 16763 |
| | D4 I | 20591 | 574962 | 17190 | 17078 | 17176 | 18047 |
| | G6 | 27180 | 213569 | 796289 | 22905 | 14470 | 21593 |
| | D4 II | 22785 | 64041 | 11803 | 339829 | 21765 | 23394 |
| | F4 | 22291 | 45029 | 11608 | 24051 | 249330 | 17521 |
| | G4 | 25794 | 70443 | 1119908 | 31474 | 149623 | 286265 |

| | |
|---|---|
| pGFP(TetA) positive control | 738296 |
| pTetR negative control | 22328 |

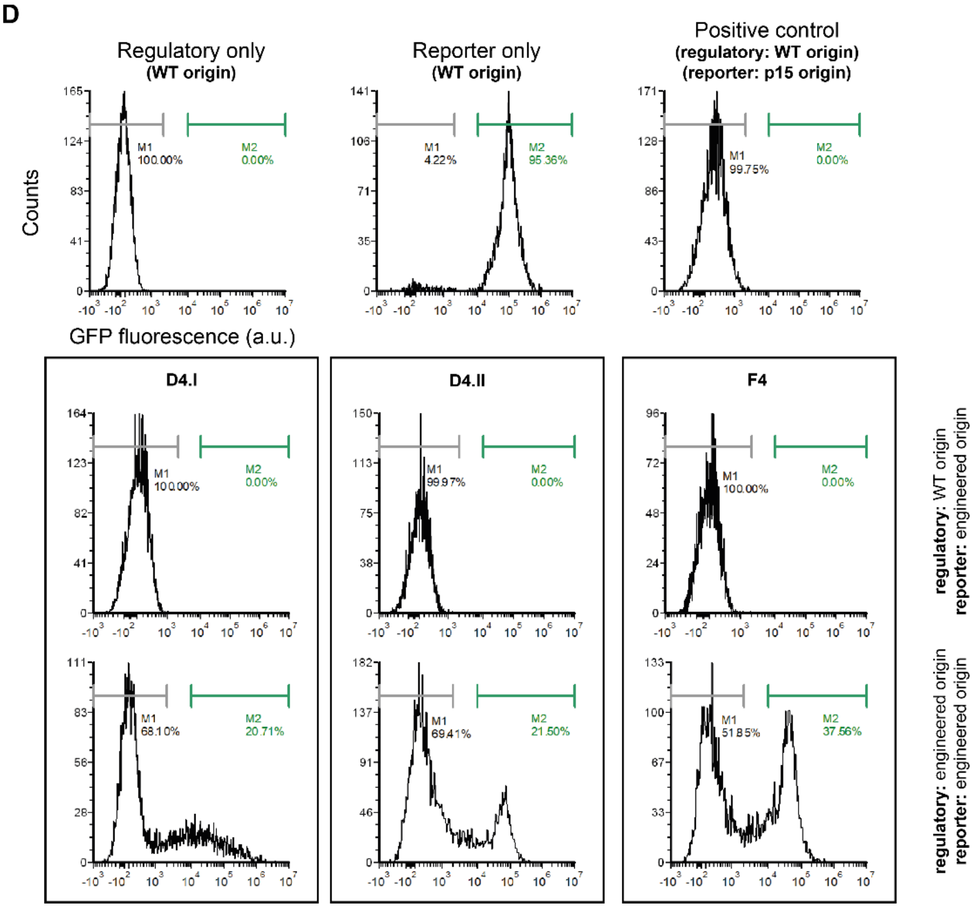

**Supplementary Figure 6: Impact of culture medium on plasmid compatibility and characterization of cross-compatibility. A.** Box plot showing the distribution of normalized fluorescence for cross-compatibility assays carried out in different culture media (single passage). Outliers (2% cut-off in ROUT analysis), that is significantly expressing GFP, are shown in red. **B.** Box plot showing the impact of passaging in cross-compatibility assays in M9 media. Outliers (2% cut-off in ROUT analysis), that is significantly expressing GFP, are shown in red. **C.** Cross-compatibility results (normalized fluorescence values) obtained in M9 after one or two passages. Outliers identified in **B.** are shown in green. The third passage is shown in Figure 4A. **D.** Flow cytometry analysis of cross-compatibility assays showing controls and selected replication origins, their compatibility to wild-type colE1 origins and their self-incompatibility. Markers show ranges used to quantify non-fluorescence (grey) and fluorescent (green) fractions of the populations. Each experiment included at least 6200 events post single-cell gating.

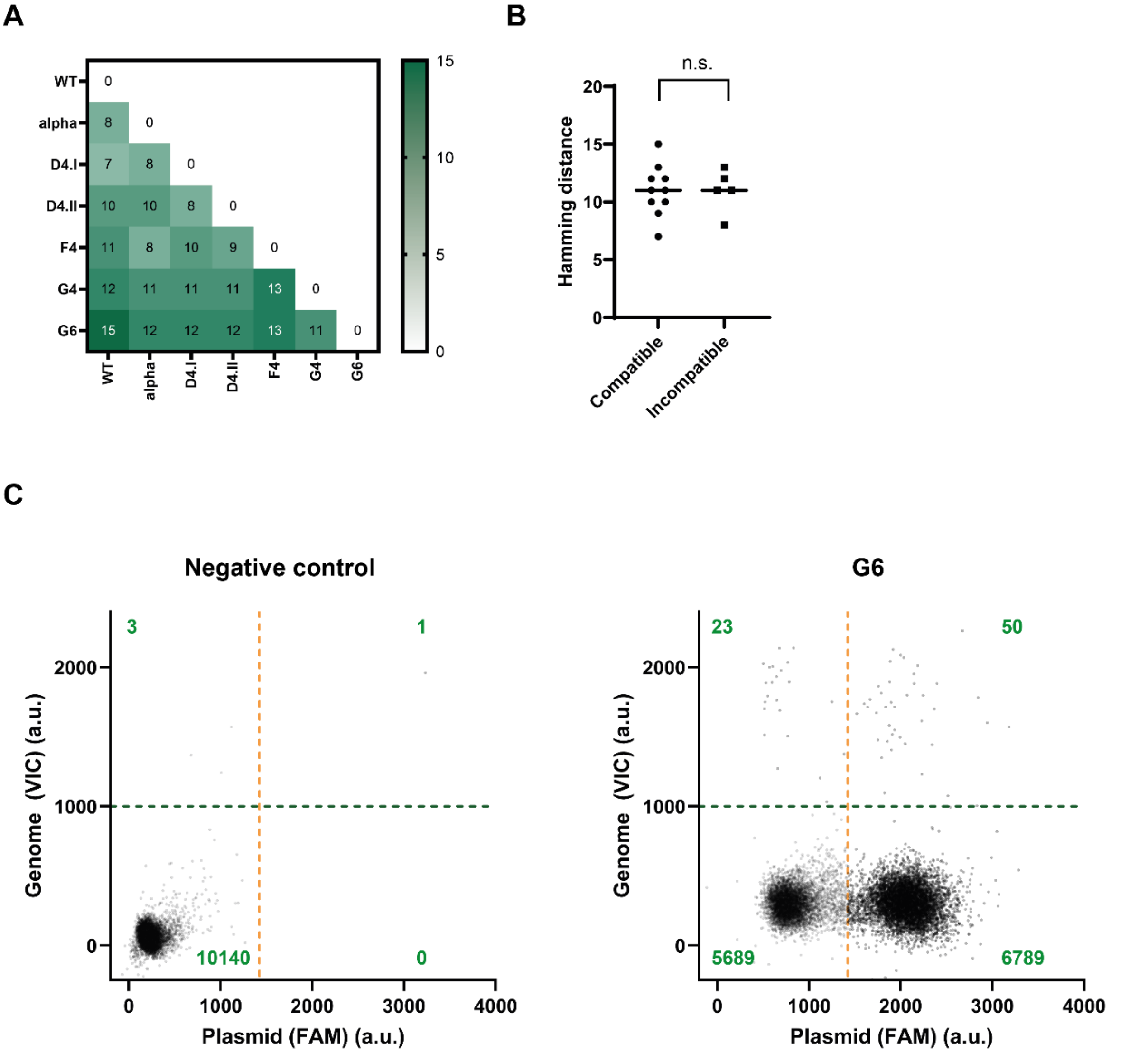

**Supplementary Figure 7: Sequence analysis of engineered colE1 origins and quantification of plasmid copy number per cell. A.** Hamming distance (number of substitutions between 2 sequences) between the engineered origins of replication. **B.** Hamming distance distribution between compatible and incompatible colE1 origins (using data presented in Figure 4A). A Kolmogorov-Smirnov test was used to compare the two Hamming distance distributions but no significant different was observed. **C.** Examples of digital PCR results to show negative control (no template) and results obtained for G6 engineered origin. The quadrants are determined automatically by the analysis program and the number of observations in each quadrant are shown in green.

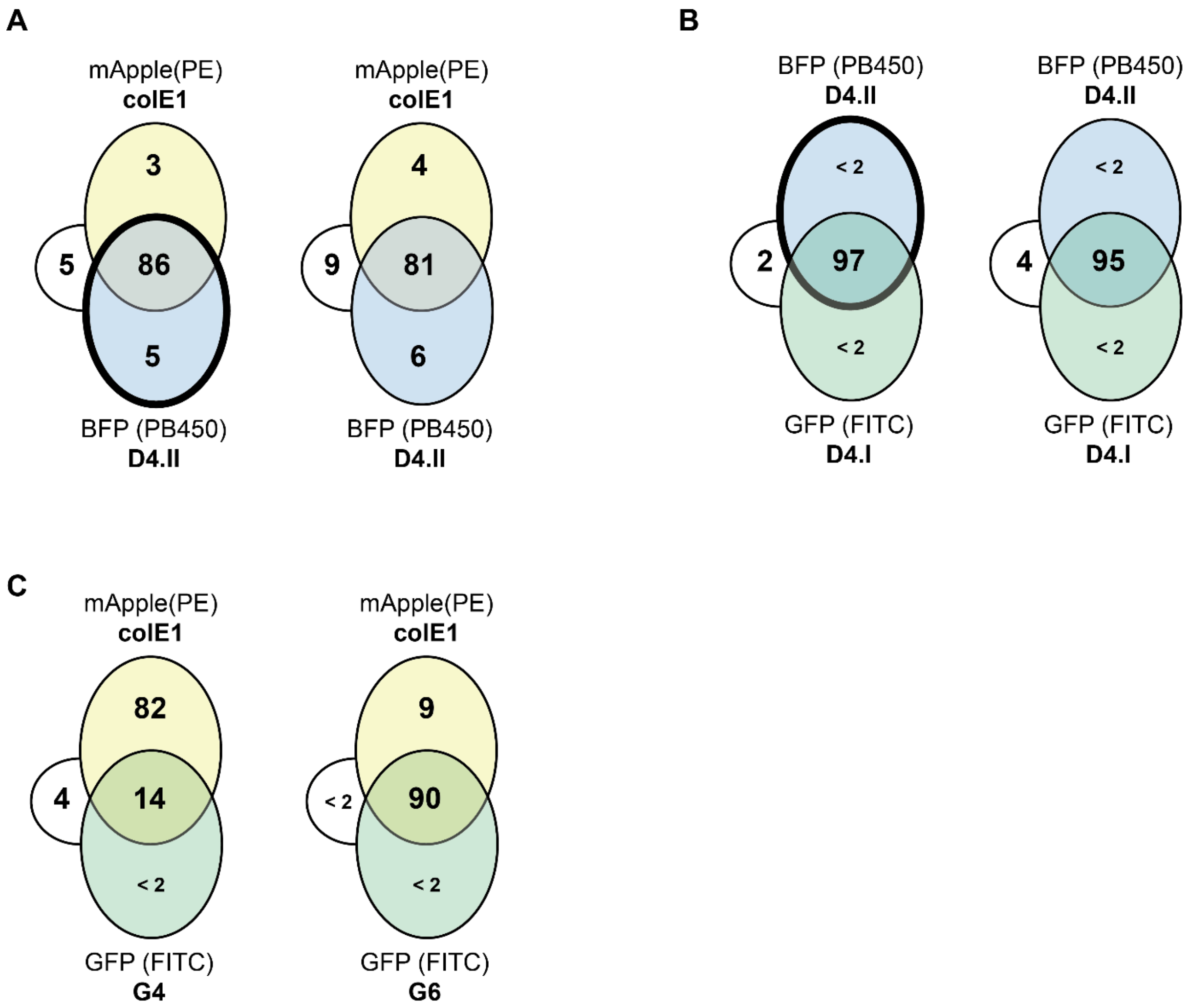

**Supplementary Figure 8: Pairwise compatibility between origins in plasmids expressing fluorescent proteins.** Summary of flow cytometry analysis of cultures post-serial passaging in M9 used to investigate plasmid retention and plasmid compatibility. Origins and fluorescent protein encoded are shown around the edges of the Venn diagram: D4.II origin in mTagBFP2-pBAD (blue), colE1 origin in mApple-pBAD (yellow) and other origins in GFP-pBAD (green). Thick borders show experiments where chloramphenicol was used to ensure D4.II plasmid retention. **A.** D4.II and colE1 origins. **B.** D4.I and D4.II origins. Both show that chloramphenicol selection has little impact on the retention of the plasmids. **C.** colE1 origins and G4 or G6. Under the culture conditions used for this experiment, the G4 origin is lost from the population (in alignment with what was seen in SI Fig 5C, but different from what was observed in the high-throughput assay (Figure 4A). These experiments were also used as controls for the 3-way intercompatibility assays.

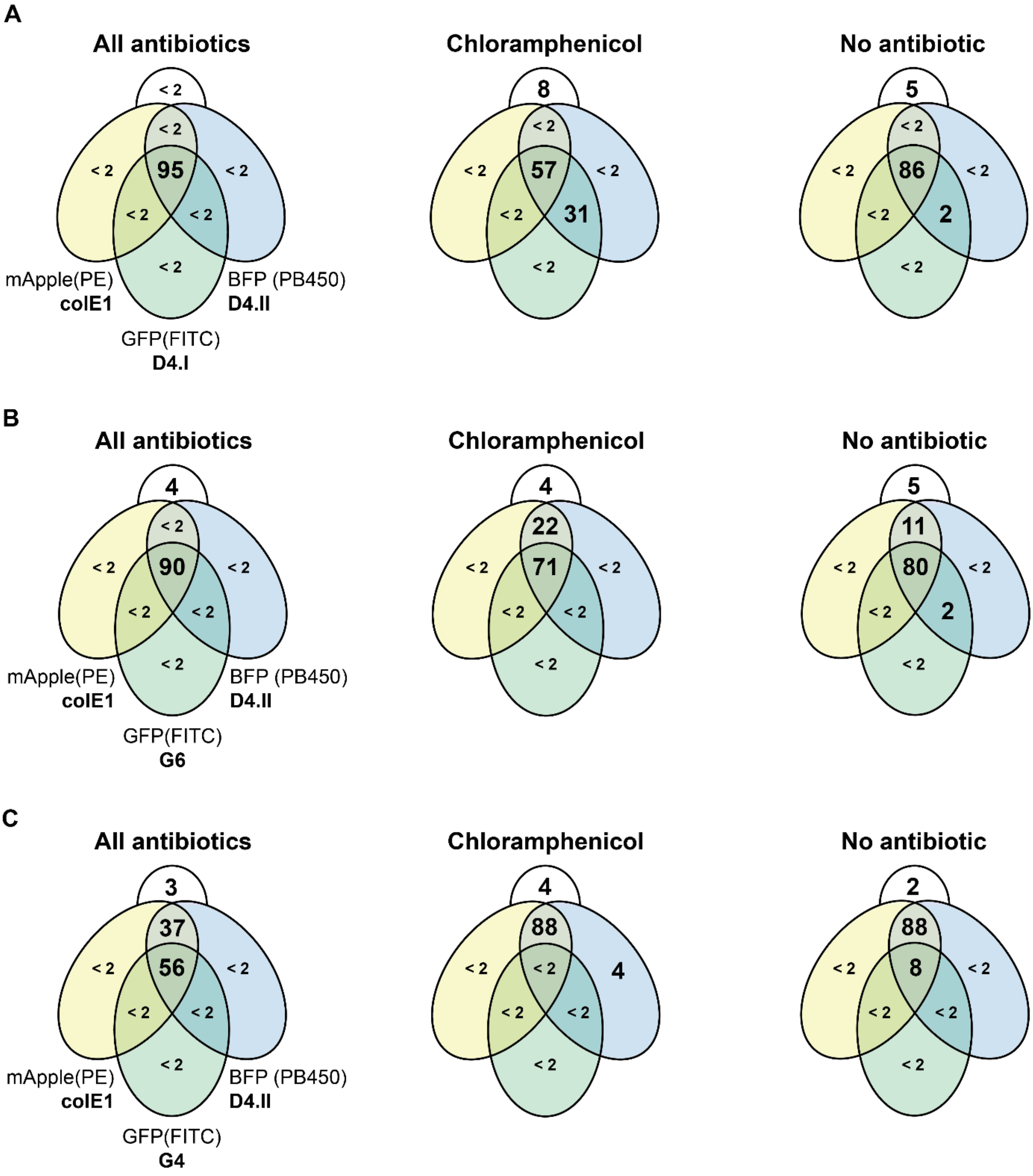

**Supplementary Figure 9: Plasmid intercompatibility assays.** Summary of flow cytometry analysis of cultures post-serial passaging in M9 used to investigate plasmid retention and plasmid compatibility. Cells co-transformed with three plasmids harbouring different plasmid origin combinations were serially passaged in M9 before being analysed by flow cytometry to determine which plasmids had been retained in culture. Plasmid origins and fluorescent proteins are shown for each combination around the Venn diagram. BFP is shown in blue, mApple in yellow and GFP in green. Cultures were maintained with all antibiotics (ampicillin, chloramphenicol and kanamycin), or with only chloramphenicol, or without any added antibiotics. **A.** Origins D4.I, D4.II and wild-type colE1. Passaging of the culture in the presence of chloramphenicol results in significant wild-type colE1 loss. **B.** Origins G6, D4.II and colE1. Plasmid harbouring G6 origin is preferentially lost from culture but at slow rates, ensuring that most of the population retains all 3 plasmids. **C.** Origins G4, D4.II and wild-type colE1. In contrast to the pairwise assays, plasmids with the G4 origin were rapidly lost from the population, even in the presence of all three antibiotics, suggesting that it may not as stable as other origins or that its low copy number puts it in a significant disadvantage during replication.

**Supplementary notes:**

**Polymerase Chain Reaction.** PCR was used to generate the biological constructs for this work. Unless stated otherwise, all reactions were carried out in 50 µL with the following reaction components: 1X Q5 reaction buffer, 0.5 µM of each primer, 200 µM dNTPs, 0.2 ng/µL of template, 0.02 U/µl Q5 enzyme (New England Biolabs), and deionized sterile water to complete the reaction volume. The reaction conditions typically consisted of an initial denaturation at 95°C for 30 seconds, followed by 30 – 32 cycles of 95°C for 20 seconds, 50 - 72°C for 30 seconds, 72°C for 30 seconds/kb of the target DNA product. All reactions included final 72°C extension for 5 minutes.

**Raw data.** All data and analyses generated in this project are publicly available at https://github.com/PinheiroLab/Engineered_colE1_origins.
